# K48- and K63-linked ubiquitin chain interactome reveals branch- and chain length-specific ubiquitin interactors

**DOI:** 10.1101/2024.01.08.574586

**Authors:** Anita Waltho, Oliver Popp, Christopher Lenz, Lukas Pluska, Volker Dötsch, Phillip Mertins, Thomas Sommer

## Abstract

The ubiquitin (Ub) code denotes the complex Ub architectures, including Ub chains of different length, linkage-type and linkage combinations, which enable ubiquitination to control a wide range of protein fates. Although many linkage-specific interactors have been described, how interactors are able to decode more complex architectures is not fully understood. We conducted a Ub interactor screen, in humans and yeast, using Ub chains of varying length, as well as, homotypic and heterotypic branched chains of the two most abundant linkage types – K48- and K63-linked Ub. We identified some of the first K48/K63 branch-specific Ub interactors, including histone ADP-ribosyltransferase PARP10/ARTD10, E3 ligase UBR4 and huntingtin-interacting protein HIP1. Furthermore, we revealed the importance of chain length by identifying interactors with a preference for Ub3 over Ub2 chains, including Ub-directed endoprotease DDI2, autophagy receptor CCDC50 and p97-adaptor FAF1. Crucially, we compared datasets collected using two common DUB inhibitors – Chloroacetamide and N-ethylmaleimide. This revealed inhibitor-dependent interactors, highlighting the importance of inhibitor consideration during pulldown studies. This dataset is a key resource for understanding how the Ub code is read.

## Introduction

Ubiquitination is a post-translational modification which regulates almost every cellular process. To achieve this, ubiquitination adds a signal onto the substrate protein, recruiting specific ubiquitin-binding proteins (UBPs) via their ubiquitin-binding domains (UBDs) to carry out a desired function. There are a wide range of UBPs and functions, for example, recruitment of DNA repair proteins to the site of DNA damage^1^, endocytosis adaptors binding to a membrane receptor to initiate its vesicular transport^2^ or recruitment of the proteasome leading to substrate degradation^3^. The building block of every ubiquitination signal is just a simple 9.6KDa protein – ubiquitin (Ub). How this small protein can control this wide array of protein fates is down to the complex chain architectures that Ubs can form, known as the Ub code^4^.

Substrate ubiquitination is initiated by monoubiquitination (mono), the covalent attachment of Ub via its C-terminal hydroxide to, most conventionally, a lysine (K) of the substrate protein. This can be followed by ubiquitination of Ub itself at one of its 7 K residues or N-terminal amide group, thus forming a Ub chain. The resulting Ub2 chain can also be described by its linkage-type, the residue through which the Ub moieties are linked, for example, K48-linked Ub2. This chain can be extended to Ub3, Ub4 and so on. Ub chains can be homotypic, meaning all Ubs in the chain are linked through the same residue, or heterotypic, in which Ubs are linked through different residues. Heterotypic chains may be mixed linkage, with alternating linkage, or branched, where a single Ub in the chain has more than one Ub attached to it^5,6^. The Ub code encompasses this diverse range of Ub architectures, based on linkage-type, chain length and homotypic or heterotypic linkage.

K48-linked Ub is the most abundant linkage type in the cell, followed by K63-linked Ub^7^. The former is a well-studied proteasomal-degradation signal^4^, and the latter is associated with pathways such as autophagy^8^, protein trafficking^9,10^ and NF-κB signalling^11^. Branched Ub chains containing both linkages, referred to as K48-/K63-linked branched Ub, are also present in the cell, making up 20% of all K63 linkages^12^. The function of this chain-type is less well-defined; current literature suggests context-specific roles for K48-/K63-linked branched Ub, in one instance enhancing NF-κB signalling^12^, and in another triggering proteasomal degradation^13^. Furthermore, K48/K63 branch-specific binders are an only recently emerging area of investigation^14,15^.

Cell-wide Ub-interactor pulldown studies enable us to decode, meaning reveal the function of, Ub signals through identification of chain type-specific UBPs. Furthermore, information on UBP specificity aids our understanding of the mechanism of Ub binding and the role of UBDs. Thus far, published datasets have used chemically synthesised Ub chains to identify potential chain linkage-^16,17^, chain length-^18^ and branched-dependent^15^ Ub interactors in humans. Our dataset builds on this information using native enzymatically synthesised Ub chains to probe for chain length- and branch-specific interactors of K48- and K63-linked Ub chains in both humans and budding yeast. We identified interactors with a preference for Ub3 over Ub2, including DDI1, yeast homologue Ddi1, CCDC50 and FAF1, and K48/K63-linked branch-specific interactors, including PARP10, UBR4 and HIP1. We were able to validate HIP1’s K48/K63-linked branched Ub preference by Surface Plasmon Resonance (SPR). Furthermore, we investigated, by comparison, the effect of reagents commonly used as DUB inhibitors, N-ethylmaleimide (NEM) and Chloroacetamide (CAA), on Ub binding.

## Results

### Ubiquitin interactor screen establishment

We designed a K48- and K63-linked Ub interactor screen in which Ub chains are immobilized on resin and used as bait to enrich Ub interactors from cell lysate. Interactors are then identified by liquid chromatography-mass spectrometry (LC-MS) and chain type enrichment patterns analysed by statistical comparison (Fig 1A). As we are interested in comparing chain-linkage, length and branch-specific Ub interactors, we sort to synthesise mono Ub, homotypic K48- and K63-linked Ub2 and Ub3 and K48/K63-linked branched Ub3. We chose to use a K48/K63-linked branched Ub3 which resembles the branchpoint, the basic unit of a branched chain, as the complex architecture of branched chains in the cell is not fully elucidated and may vary in different contexts. We previously discovered the K48-branching activity of the E2 Ub-conjugating enzyme Ubc1^19^ , thus using this enzyme, along with K48- and K63-specific E2 enzymes, CDC34 and Ubc13/Uev1a, we were able to enzymatically synthesise and purify our desired Ub chains *in vitro.* Chain linkage composition was confirmed using the UbiCrest method^20^ by selective disassembly with the K48- and K63-specific deubiquitinases (DUBs) OTUB1 and AMSH, respectively (S.Fig 1A).

**Figure 1:**
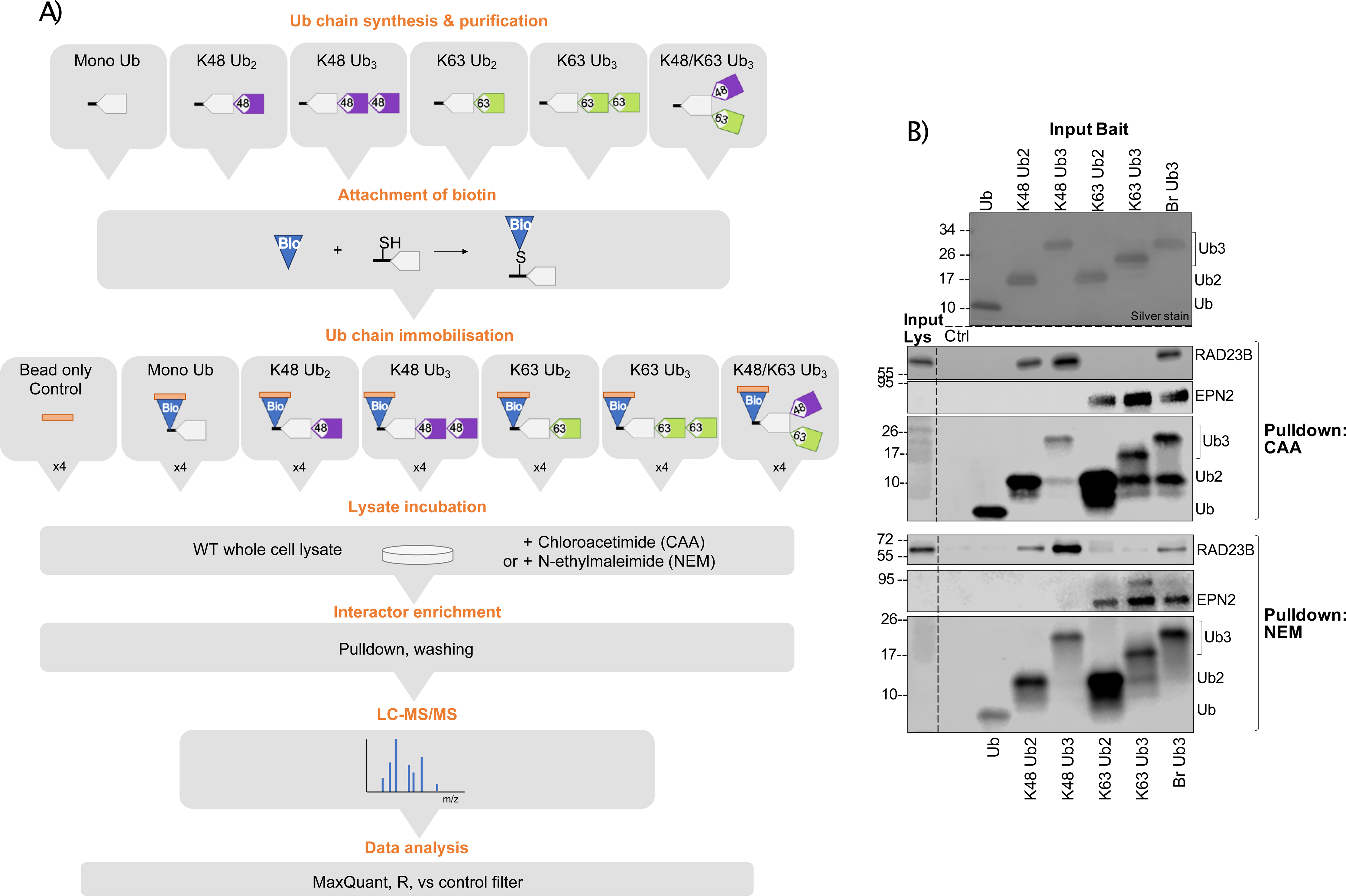
Ubiquitin chain interactor pulldown using different DUB inhibitors. A) Schematic of Ub interactor pulldown and MS experiment. B) Western blot of Ub interactor pulldown using different cysteine alkylators, CAA and NEM, as DUB inhibitors. Silver stain of input Ub. Pulldown blotted using anti-RAD23B, anti-EPN2 and anti-Ub antibodies.

In order to immobilize the Ub chains on streptavidin resin, we inserted a cysteine residue near the C-terminus of the proximal Ub of each chain and attached a biotin molecule specifically using a cysteine-maleimide reaction. Complete biotin conjugation was confirmed using intact MS (S.Fig 1B-G). These Ub chains contain native isopeptide bonds, therefore they are susceptible to chain disassembly by endogenous DUBs in the lysate. As cysteine proteases are the largest DUB family, cysteine alkylators including Chloroacetamide (CAA) and N-Ethylmaleimide (NEM) are often used as DUB inhibitors ^20–23^ . However, cysteine alkylators don’t just target DUBs, they can theoretically alkylate any exposed cysteine on a protein.

Furthermore, whilst CAA is relatively cysteine-specific^24^, NEM, when used for peptide alkylation for MS, was found to have frequent side reactions with N-termini and lysine side chains^25^. Off-target effects on non-DUB proteins are a concern as they could alter Ub binding surfaces. As an example, NEM combined with Iodoacetamide (IAA) treatment was found to perturb NEMO binding to K63-linked Ub chains *in vitro*^26^. We tested CAA and NEM for their ability to stabilize immobilized Ub chains in HeLa cell lysate (Fig 1B). Note that, the anti-Ub antibody showed some linkage-dependent binding (S.Fig 10A). Known linkage-specific UBPs, K48-specific UBP RAD23B^27^ and K63-specific UBP EPN2^17^ were used as controls. With either CAA or NEM, RAD23B and EPN2 were only enriched on their preferred linkage-types showing that both inhibitors block chain disassembly sufficiently for specific UBP pulldown. However, there were differences in the stability of immobilized Ub in each inhibitor treated lysate; in NEM there was nearly no chain disassembly, whereas in CAA Ub3 was partially disassembled to Ub2 (Fig 1B). This could be expected as NEM is a more potent cysteine alkylator ^21^. Whilst, acknowledging the limitation of partial digestion under CAA treatment, it is notable that the original bait remains the predominant Ub species throughout the experiment. Moreover, the Ub bait abundance significantly exceeds that of the UBPs in the lysate. Taken together with the distinct binding patterns of the known UBPs, we suggest that the CAA approach is effective in selectively enriching chain-specific UBPs, despite partial chain digest. In summation of the discussed advantages and disadvantages of using either CAA or NEM, we chose to perform the Ub interactor screen with each inhibitor separately, so that the comparison of datasets can reveal both overlapping and inhibitor-specific Ub interactors.

### Ubiquitin interactor enrichment patterns

Using our established set up, we identified Ub interactors from inhibitor-treated HeLa and *S. cerevisiae* lysate (Fig 1A). Principle component analysis showed clustering of samples by bait type (S.Fig 2A and B.). After filtering, normalization and imputation, 4540 and 4526 unique protein isoforms were identified across pulldowns from CAA- and NEM-treated HeLa lysate, respectively, with an overlap of 3711. For the CAA dataset, this included 215 expected UBPs, according to a protein list made from collating proteins found under the Gene Ontology term Ub-binding (0043130) and UBD-containing proteins from the UUICD database. The NEM dataset included 195 expected UBPs (S.Fig 3A). To remove background binders and interrogate chain-type dependent enrichment patterns, proteins were prefiltered by significant enrichment in any Ub chain pulldown in comparison to the bead-only control (two-sample moderated T-test, Adj.P < 0.05). This increased the proportion of expected UBPs to 104/544 and 64/206 prefiltered protein isoforms in CAA and NEM datasets, respectively (S.Fig 3B), suggesting that our prefiltering method positively selects for UBPs.

To identify chain type-specific enrichment patterns, we compared interactomes across all Ub chain types generating 286 and 139 significant differently enriched proteins in CAA and NEM datasets, respectively (moderated F test, Adj.P < 0.05). This included 81 expected UBPs with CAA and 54 with NEM (Fig 2A). Comparison to a published HeLa global absolute proteome^28^ revealed that our significant interactors are not biased towards highly abundant proteins (S.Fig 2). Significant interactors for each chain type were well correlated between NEM and CAA datasets, especially for K48/K63-branched chain interactors (Fig 2B). In order to identify inhibitor-specific binding patterns, we compared the enrichment across chain types of each significant hit in both datasets (S.Fig 4). A number of 53 significant hits in the CAA dataset were not identified in any pulldown with NEM-treated lysate, including 7 expected UBPs, and 13 significant hits in the NEM dataset were not identified with CAA (S.Fig 4C, E and F). Most significant hits which were also expected UBPs had the same enrichment patterns across datasets (S.Fig 4A). However, interestingly, IKBKG/NEMO, an expected K63-specific UBP, was significantly enriched on K63-linked chains in CAA and on K48-linked chains in NEM, an effect that was previously seen *in vitro*^26^. Several other expected UBPs with significant preference for K63-linked Ub with CAA, had more general preference for both K48- and K63-linked triUb with NEM, including MINDY3, BIRC2, GGA3, DNAJB2, XIAP and TSG101. A similar pattern of increased enrichment on K48-linked Ub3 with NEM-treated lysate was also seen for several significant hits that are not expected UBPs: KATNAL2, STON2, UBFD1, NIPSNAP2, LACTB and NIPSNAP1 (S.Fig 4B). These observations may be a result of increased chain stability by the more potent DUB inhibitor NEM or unspecific alkylation affecting Ub-binding sites, as is the case for IKBKG/NEMO^26^.

**Figure 2:**
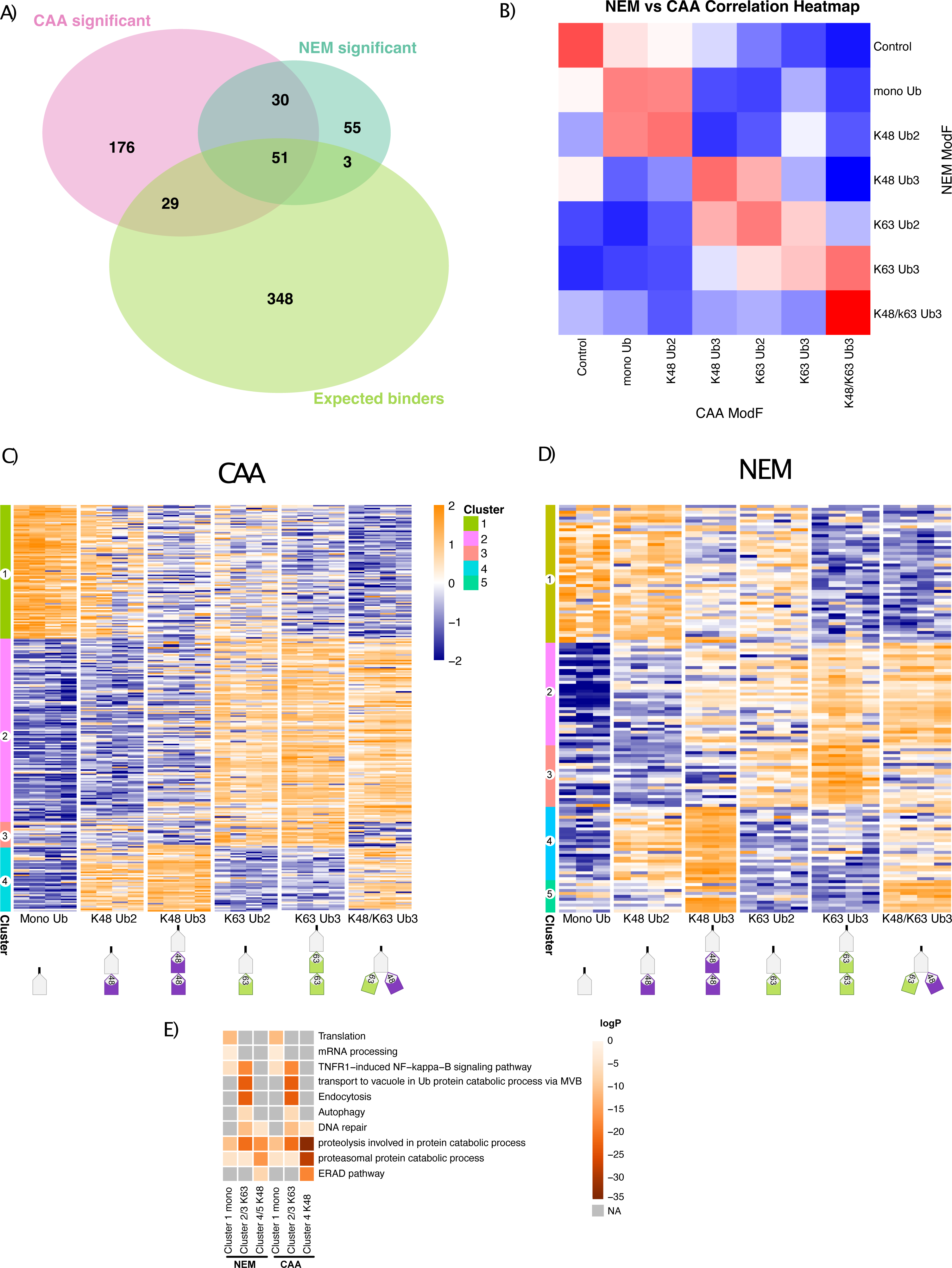
Conserved interactor enrichment patterns between different DUB inhibitor datasets. A) Overlap of significant differently enriched proteins across Ub pull-downs from CAA- and NEM-treated lysate with expected Ub binders. Significant hits identified by moderated F test (adj.P cutoff < 0.05). Data was prefiltered Ub enrichment (two-sample moderated t-test comparing experimental groups with the bead control, logFC > 0, adj.P cutoff < 0.05). Expected binders list compiled from Gene Ontology term Ub-binding 0043130 and UBD-containing proteins from the IUUCD bioCuckoo database. B) Correlation of Ub chain interactomes between CAA and NEM datasets. Spearman correlation calculated by comparing the moderated F value for each pulldown between datasets. No P value cut off used. C and D) Clustering of significant differently enriched proteins from (C) CAA and (D) NEM datasets. Significance as in A. Hierarchical clustering by Euclidean distance. Heatmap of iBAQ values, scaled by row using z scoring. E) Heatmap of Gene Ontology (GO) enrichment of clusters. GO enrichment calculated with Metascape, with a minimum overlap of 2, P value cut off < 0.01 and minimum enrichment of 1.5. Grey is not enriched. Some GO terms are abbreviated for the figure: TNFR1−induced NF−kappa−B is TNFR1−induced NF−kappa−B signalling pathway and transport to vacuole in Ub protein catabolic process via MVB is protein transport to vacuole involved in ubiquitin−dependent protein catabolic process via the multivesicular body sorting pathway

Significant proteins clustered into 4 or 5 similar clusters for CAA and NEM, respectively: proteins significantly enriched on mono Ub and Ub2 (Cluster 1), on K63-linked Ub (Cluster 2 and 3) and on K48-linked Ub (Cluster 4 and 5) (Fig 2 C and D, more detailed in S.Fig 5 and 6). In Cluster 1, there is an overlap of only 4 proteins between datasets: known Ub-binding autophagy receptor SQSTM1/p62, mitochondrial matrix protein NIPSNAP2, arginine methyl transferase PRMT5 and mitochondrial inner membrane space serine protease LACTB (S.Fig 7A). Interestingly, NIPSNAP2 and SQSTM1/p62 have been reported to interact with each other during the initial stages of mitophagy^29^. Gene ontology (GO) enrichment analysis revealed an enrichment of translation and mRNA processing -related proteins in Cluster 1 across datasets (Fig 2.E).

There were 42 proteins in common between datasets with a binding preference for K63-linked Ub (Cluster 2 and 3), including known K63-specific UBPs: Endosomal Sorting Complex Required for Transport (ESCRT) proteins STAM^30^, STAM2 and TOM1^31^, endocytic adaptor proteins ANKRD13A, ANKRD13B and ANKRD13D^32^, BRCA1-A subunit UIMC1^33,34^, IL-1 signalling-related protein TAB2^35^, and branching E3 ligase HUWE1^12^ (S.Fig 7 D). Interestingly, also in Cluster 2 is K48-processing DUB MINDY3, which was recently found to prefer cleaving K48-linked Ub within a branched (K48)/K63 Ub4 chain^14^. Proteins in Clusters 2 and 3 were associated with NF-κB signalling, Ub-dependent vacuolar transport, endocytosis and autophagy (Fig 2E), in line with current literature on K63-linked chains^8–11^. Interestingly, whilst most proteins interacted with K63 linkages in both homotypic and branched heterotypic chains to a relatively equal extent (Cluster 2)(S.Fig 7B), there was also a cluster of K63-linked Ub-specific proteins which were less enriched on branched chains (Cluster 3), including, conserved across datasets, ESCRT component Tom1 (S.Fig 7C).

Proteins enriched on K48-linked Ub (Cluster 4 in the CAA dataset and Cluster 4 and 5 in the NEM dataset) included known K48-specific UBPs: proteasomal shuttle factors RAD23A^36^ and RAD23B^27^, VCP/p97 adaptors UFD1^37^ and FAF1^38^, and DUBs ATXN3^34,39^, OTUD5^40^ and MINDY1^41^ (S.Fig 7 E). As expected from the literature, proteins in these clusters were strongly associated with proteasomal degradation and endoplasmic reticulum-associated degradation (ERAD)^4^ (Fig 2E). Interestingly, in the NEM dataset, interactors enriched on K48-linked Ub were separated into two clusters: proteins with a preference for homotypic K48-linked Ub3 (Cluster 4) and those with a branched K48/K63-linked Ub3 preference or equally enriched on both homotypic and branched Ub3 (Cluster 5) (S.Fig 6). The former (Cluster 4) includes proteasome regulatory subunit PSMD4, proteasomal shuttle factors RAD23A and RAD23B, DUB MINDY2 and VCP/p97 adaptor UBXN1. The latter (Cluster 5) includes DUB MINDY1 which has selectivity for long K48-linked chains^42^, but was recently found to preferentially cleave heterotypic K48/K63-linked chains^14^. Taken together, the clustering results support data quality as the chain preference of known linkage-specific UBPs and enrichment of linkage-specific pathways is reproduced. This heatmap also provides Ub linkage-specific binding patterns for potential novel UBPs or UBPs whose chain preference was previously unknown.

Our yeast Ub interactor screen identified 2315 unique protein isoforms, after filtering, normalization and imputation, including 76 expected UBPs (S.Fig 3C). Prefiltering for Ub-enriched proteins, by significant Ub preference over the control as described above, resulted in 247 proteins, including 38 expected UBPs (two-sample moderated T-test, Adj.P < 0.05)(S.Fig 3D). We compared interactomes across Ub chain types generating 148 significant differently enriched proteins (moderated F test, Adj.P < 0.05) (S.Fig 3E). Significant proteins clustered by into 3 clusters: proteins significantly enriched on mono Ub and partially on K63-linked Ub2 (Cluster 1), on K63-linked Ub (Cluster 2) or on K48-linked Ub (Cluster 3) (S.Fig 8). Cluster 1 included expected UBPs with UBDs, but whose chain-type specificity was unknown, for example, Prp-containing Duf1 and UBA-containing Gts1^43,44^.

Amongst interactors enriched on K63-linked Ub (Cluster 2) were known K63-specific UBPs including endocytic regulator Ent2^45^, clathrin adaptor Gga2^45^, ESCRT components Vps27and Hse1 ^45^. Cluster 2 also contained UBPs with known UBDs, but unknown chain-type preference, including LSB5 which contains GAT and VHS domains, and Rsp5 cofactor Rup1 which has a UBA domain^46^. K48-linked Ub enriched proteins (Cluster 3) include known K48-specific UBPs, for example, proteasomal receptor Rpn10^45^, Cdc48 adaptor proteins Npl4^47^ and Shp1^45^, and proteasomal shuttle protein Rad23^45^. Expected UBP Cia1, which has a Prp UBD and for which chain type specificity was previously unknown, was also in Cluster 3. Also, in the K48-linked Ub enriched cluster was C-terminal hydrolase DUB Yuh1. We validated Yuh1 K48-linked Ub preference by Western Blot (S.Fig 10C).

Proteasomal subunits Rpt1, Rpt2, Rpt3, Rpt4, Rpt5, Rpt6, Rpn2, Rpn3, Rpn5, Rpn6, Rpn7, Rpn8, Rpn9, Rpn12 and Cdc48 and its adaptors Ufd1 and Ubx5 were enriched on K48-linked Ub (Cluster 3), although they have been shown to not directly bind to Ub^45,48,49^ (S.Fig 8). It must, therefore, be taken into consideration that some proteins enriched on Ub, by this method, are part of larger Ub-binding complexes, rather than direct UBPs themselves. We validated the chain specific pulldown of several known UBPs: Vps9, Dsk2, Rad23, Ddi1 and Npl4 by Western Blot (S.Fig 10C). Overall, the yeast dataset successfully reproduced the chain preference of known linkage-specific UBPs, whilst it also provided chain type-specificity information for UBPs with previously unknown preference.

### Length-dependent Ub interactors

Not only linkage-type, but also length of Ub chain determines interactor binding. Previous Ub interactor MS screens have identified UBPs which only interact with long M1-linked^17^, K27-linked, K29-linked or K33-linked chains ^18^. Some proteins bind to multiple Ub moieties simultaneously, including ZFAND2B/AIRAPL and DUB USP5/IsoT which have 3 or 4 Ub binding sites, respectively^50,51^. Furthermore, it is conventionally believed that the proteasome requires K48-linked ≥Ub4^52,53^, although other papers contest this^54,55^. DUBs have also been shown to have chain length preference, for example MINDY1 prefers longer chains^56,57^ and UCHL3 prefers shorter chains^58^. Cell ubiquitomes also reveal chain length information; in yeast almost 50% of Ub detected existed in monomers and the average chain length varied depending on the linkage type^59^.

With these findings in mind, we set out to compare the interactomes of homotypic K48- and K63-linked Ub2 and Ub3 chains in our data. In both the CAA and NEM dataset, there was a stronger correlation between K63-linked Ub2 and Ub3 interactomes, than between K48-linked Ub2 and Ub3 (Fig 3A and B). This observation was more apparent in the NEM dataset compared to the CAA dataset. Furthermore, with NEM there were 99 and 13 significant differently enriched proteins in the Ub3 versus Ub2 comparison for K48- and K63-linked Ub, respectively, compared to only 20 and 10 for CAA (moderated T-test, Adj.P < 0.05). This could be the result of Ub3 disassembly to Ub2 in the CAA-treated lysate pulldown (Fig 1B). Interestingly, in both datasets the monoUb interactome is the most well-correlated with the K48-linked Ub2 interactome (S.Fig 9A and B). Taken together, these results suggest that the effect of chain length on UBP specificity varies depending on the linkage type.

**Figure 3:**
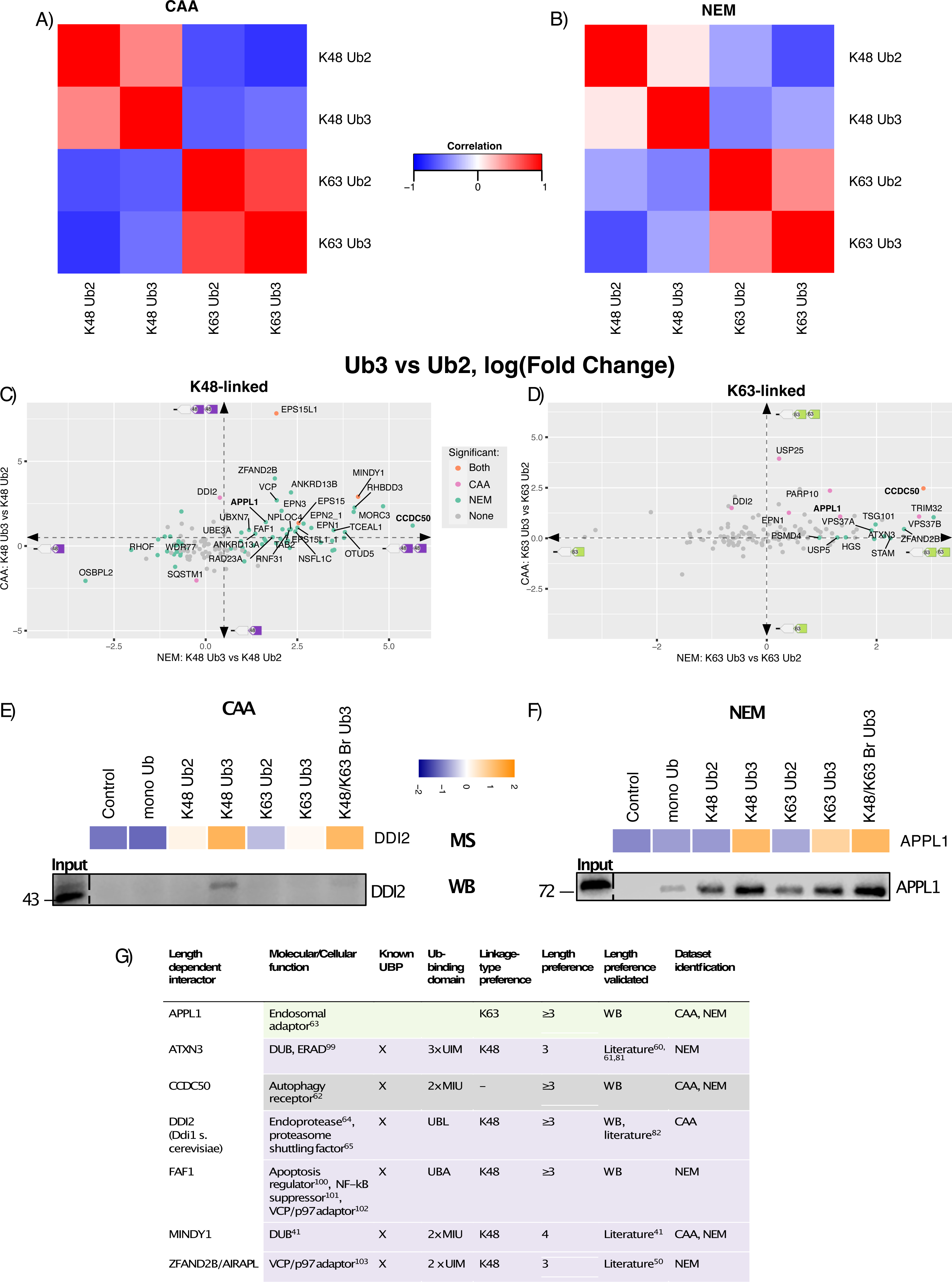
Ubiquitin chain length-dependent interactors identified. A and B) Correlation of Ub chain interactomes within (A) CAA and (B) NEM datasets. Spearman correlation calculated by comparing the moderated F value for each Ub pulldown of significant hits within each dataset. Significant hits identified as in Fig 2A (moderated F test of prefiltered Ub enriched proteins, adj.P cut off < 0.05). C and D) Scatterplot of Ub2 versus Ub3 interactor comparisons for (C) K48-linked and (D) K63-linked Ub chains across CAA (y axis) and NEM (x axis) datasets. Comparisons are made by two sample moderated T-tests of Ub3 versus Ub2 interactomes of each linkage type. Dot colours: orange is statistically significant in both datasets, pink is significant in CAA dataset only, teal blue is significant in NEM dataset only and grey is not significant in either dataset. Labelled proteins are significant in either dataset and have the same Fold Change tendencies in both datasets, log(Fold Change) < -0.5 or > 0.5 (for K48) or < 0 or > 0 (for K63) (as shown by dotted arrow lines). DDI2 is labelled despite NEM log(FC) = 0.39 due to interest based on same length preference of yeast homologue Ddi1. Adj.P cut off < 0.05. E and F) MS heatmap and Western blot of pulldown of length dependent Ub binders, DDI2 (E) and APPL1 (F) from CAA-treated and NEM-treated lysate, respectively. Heatmap of modF values (Adj.P < 0.05). Same Ub interactor pulldown experiment as in Figure 1 B, thus same silver stain of input Ub. Pulldown blotted using anti-APPL1 and anti-DD2 antibodies. G) Table of identified Ub chain length dependent interactors. Linkage type preference identified from two sample moderated T-test of K48-linked Ub3 versus K63-linked Ub3. Adj.P cut off < 0.05.

3 proteins were significantly enriched on K48-linked Ub3 over Ub2 in both datasets: ESP15, ESP15L1 and MINDY1, and 27 were significant in NEM only, but with the same chain-length preference in CAA (two-sample moderated T-test, log(FC) > 0.5, Adj.P < 0.05) (Fig 3C and G). Multiple known K48-specific UBPs were amongst these, including RAD23A^36^, FAF1^38^, ZFAND2B^50^, and DUBs ATXN3^34^, MINDY1^41^ and OTUD5^40^. ZFAND2B, ATXN3 and MINDY1 were also previously shown to bind longer chains ^41,50,60,61^, thus supporting our findings. Interestingly, some of the interactors with preference for the K48-linked Ub3 over Ub2 chain are known K63-specific UBPs, including ANKRD13A, ANKRD13B^32^, EPN1, EPN2^17^, and TAB2^35^. This phenomenon is in line with a previous finding in which a K63-specific UBD also bound longer K48-linked Ub chains by avid binding to non-adjacent Ub moieties^34^. Additionally, it has been observed in yeast that some K63-specific UBPs can bind longer K48-linked chains^45^. In the same pairwise comparison, 4 proteins were significantly enriched on K48-linked Ub2 over Ub3 in the NEM dataset: SQSTM1/p62, OSBPL2, RHOF1 and WDR77, with the same enrichment pattern in CAA, but above the significance cutoff (two-sample moderated T-test, log(FC) < 0.5, Adj.P < 0.05) (Fig 3C).

In the K63-linked Ub pairwise comparison, there was only 1 protein which was significantly enriched on Ub3 compared to Ub2 in both datasets: autophagy receptor CCDC50^62^, 2 significant in NEM only: ESCRT-I components TSG101 and VPS37B, and 3 significant in CAA only: endosomal adaptor APPL1^63^, E3 ligase TRIM32 and ADP-ribosyltransferase PARP10, however each had chain-length preference conserved between datasets (two sample moderated T-test, logFC > 0.5, AdjP < 0.05) (Fig 3D and G). In the CAA dataset only, the Ub-directed endoprotease DDI2^64^ had significant preference for both K48- and K63-linked Ub3, compared to their Ub2 counterparts (Fig 3C and D, S.Fig 9A). By Ub interactor pulldown with Western Blot, we were able to validate DDI2’s preference for K48-linked Ub3 over Ub2 in CAA-treated lysate and APPL1’s preference for K48- and K63-linked Ub3 over Ub2 in NEM-treated lysate (Fig. 3 E and F). In budding yeast the only interactors enriched on Ub3 over Ub2 independent of linkage type were, the yeast homologue of DDI2, Ddi1 and Pmt4 (S.Fig 9C). We validated the Ub3 preference of Ddi1 by pulldown and Western Blot (S.Fig 10C).

Next, we sort to expand our length preference study with Ub4. Difficultly equalizing the immobilized Ub inputs and Ub4 disassembly in the lysate led to less Ub4 than other chain types in the Ub interactor pulldown (S.Fig 11). Thus, it is difficult to elucidate Ub4 binding preference as interactors may appear less enriched on Ub4 for these reasons. Nevertheless, we were able to observe K48-linked Ub3 over Ub2 preference for FAF1 and DDI2, and K63-linked Ub3 over Ub2 preference for CDCC50 with either inhibitor. Again, general Ub3 over Ub2 preference for APPL1 was only observed with NEM lysate (S.Fig 11B). In summary, comparison of Ub2 versus Ub3 interactomes revealed that the effect of chain length on interactor binding is linkage-dependent, with more interactors significantly enriched on Ub3 over Ub2 in K48-linked comparisons, than K63-linked.

### Branched chain interactors

In this screen we also investigated cell-wide branched Ub chain interactors. The K48/K63-linked branchpoint contains three Ub moieties and three isopeptide bonds like a homotypic Ub3, however it only contains one isopeptide bond of each linkage type like a homotypic Ub2. For this reason, we chose to compare the branched K48/K63-linked Ub3 with both homotypic K48-linked and K63-linked di and Ub3.

In yeast the branched K48/K63-linked Ub interactome correlated most with the K48-linked Ub3 interactome, whereas in the HeLa CAA dataset it correlates most with K63-linked Ub3. In the HeLa NEM dataset, the correlation between branched interactome and K48- or K63-linked Ub3 interactomes was relatively comparable. (S.Fig 9A-C).

Taken together, enrichment comparisons across Ub and pairwise comparison to homotypic chains show that the branched chain shares many interactors with each of its constitutive linkage types, thus conferring both K48-linked and K63-linked specificity (Fig 2C and D, and Fig 4A-D). This suggests it can act as a combination of both Ub signals.

**Figure 4:**
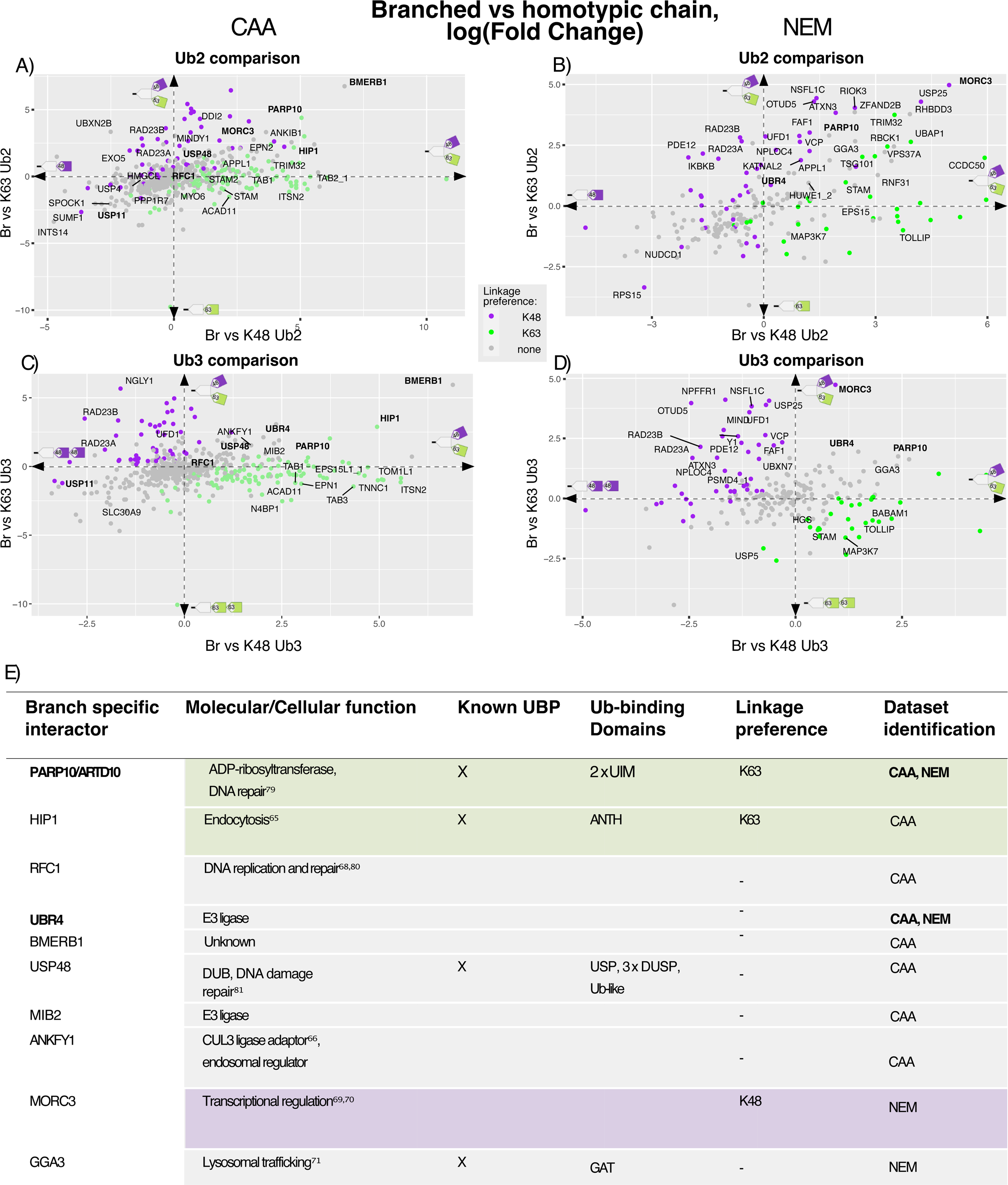
K48/K63-linked branched chain interactors identified. A-D) Scatterplots of branched versus homotypic Ub2 (A and B) or Ub3 (C and D) chain interactors in (A and C) CAA and (B and D) NEM datasets. Two sample moderated T-tests of the interactomes of K48/K63-linked branched Ub3 versus K63-linked chains, and K48/K63-linked branched Ub3 versus K48-linked Ub chains. Dot colours refer to linkage preference by two sample moderated T-test of K48-linked Ub3 versus K63-linked Ub3: purple is K48 significant, green is K63 significant and grey is not statistically significant in this comparison. Labelled proteins are significant in both branched versus homotypic chain comparisons. Proteins in bold are significant in at least two of the branched versus homotypic chain comparisons across datasets. Adj.P cut off < 0.05. E) Table of K48/K63-linked branched Ub enriched proteins, identified from comparison to homotypic Ub3 chains in C and D.

Interactors with significantly enriched on Ub3 over Ub2 homotypic chains (Fig 3G), were also a significantly enriched on branched Ub3 over homotypic Ub2, including DDI2, APPL1, FAF1, CCDC50, ATXN3 and MINDY1 (two-sample moderated T-test, Adj.P < 0.05) (Fig 4A and B). In the NEM dataset, some known K48-specific UBPs, including FAF1^38^, UFD1^37^ and ZFAND2B^50^ and K63-specific UBPs, including STAM^30^ and HUWE1^12^, had a preference for K48/K63-linked branched Ub3 over both homotypic Ub2 chains. Whereas, in the Ub3 comparison known linkage-specific UBPs were enriched on the homotypic Ub3 over branched Ub3 (Fig 4C and D).

Pairwise comparison of K48/K63-linked branched Ub to both homotypic Ub3 and Ub2 revealed two proteins with significant branched chain binding preference across datasets, ADP-ribosyltransferase PARP10 and E3 ligase UBR4 (Fig 4C-E). PARP10 contains two UIMs and was previously found to bind K63-linked ≥Ub4^65^ or K48-linked Ub2^17^. UBR4 plays a role in the formation of K11/K48-linked^66,67^ and K48/K63-linked branched Ub chains^13^. In the CAA dataset, endocytosis regulator HIP1^68^, CUL3 ligase adaptor ANKFY1^69^, DUB USP48, E3 ligase MIB2^70^, DNA polymerase accessory factor RFC1^71^ and protein of unknown function BMERB1 were additional significant branched chain enriched interactors in comparison to homotypic Ub3. Whilst with NEM, we identified transcriptional regulator MORC3^72,73^ and lysosomal trafficking factor GGA3^74^ (Fig 4C-E). Notably, RFC1 and MORC3 were also identified as branched K48/K63-linked Ub chain-specific interactors in a recent preprint^14^.

We also identified proteins with preference for homotypic chains compared to the branched chain. Both RAD23A and RAD23B were consistently significantly enriched on homotypic K48-linked chains over the branched chain. In NEM, TAB1, ACAD11 and ITSN2, and in CAA, TOLLIP and MAP3K7 were significantly enriched on homotypic K63-linked chains over the branched chains (Figure 3 A-D). Additionally, the DUB USP11 was significantly enriched on all Ub chains over the branched K48/K63-linked Ub in the CAA dataset (Fig 3 A and C). This was validated by Western Blot (S.Fig 10B).

We chose to further investigate two of the interactors enriched on K48/K63-linked branched Ub: PARP10, which was conserved across datasets, and HIP1, which was only found with CAA treatment. Ub chain pulldown with Western Blot validated our MS results (Fig 5A and B). In NEM-treated lysate, HIP1 was not sufficiently enriched in any Ub or control pulldown in line with our MS data. For PARP10 we also included Ub4 chains in the Western Blot pulldown, as literature suggests that PARP10 could bind K63-linked ≥Ub4^65^. To further validate HIP1 branched chain binding, we conducted surface plasmon resonance with different Ub chains and calculated K_D_ values using the steady state affinity model (Fig 5 C and D, and S.Fig 12). We determined a K_D_ value of 0.07µM for branched K48/K63-linked Ub3 compared to 1.55µM for K63-linked Ub3, 1.89µM for K63-linked Ub2 and 196.73µM for mono Ub. For K48-linked Ub2 and Ub3 we predict K_D_ values of 137.47 µM and 66.2 µM, respectively, however the binding affinity was outside of the concentration range tested, therefore these values may be less accurate. These results validate Hip1 as a UBP with binding preference for K48/K63-linked branched Ub.

**Figure 5:**
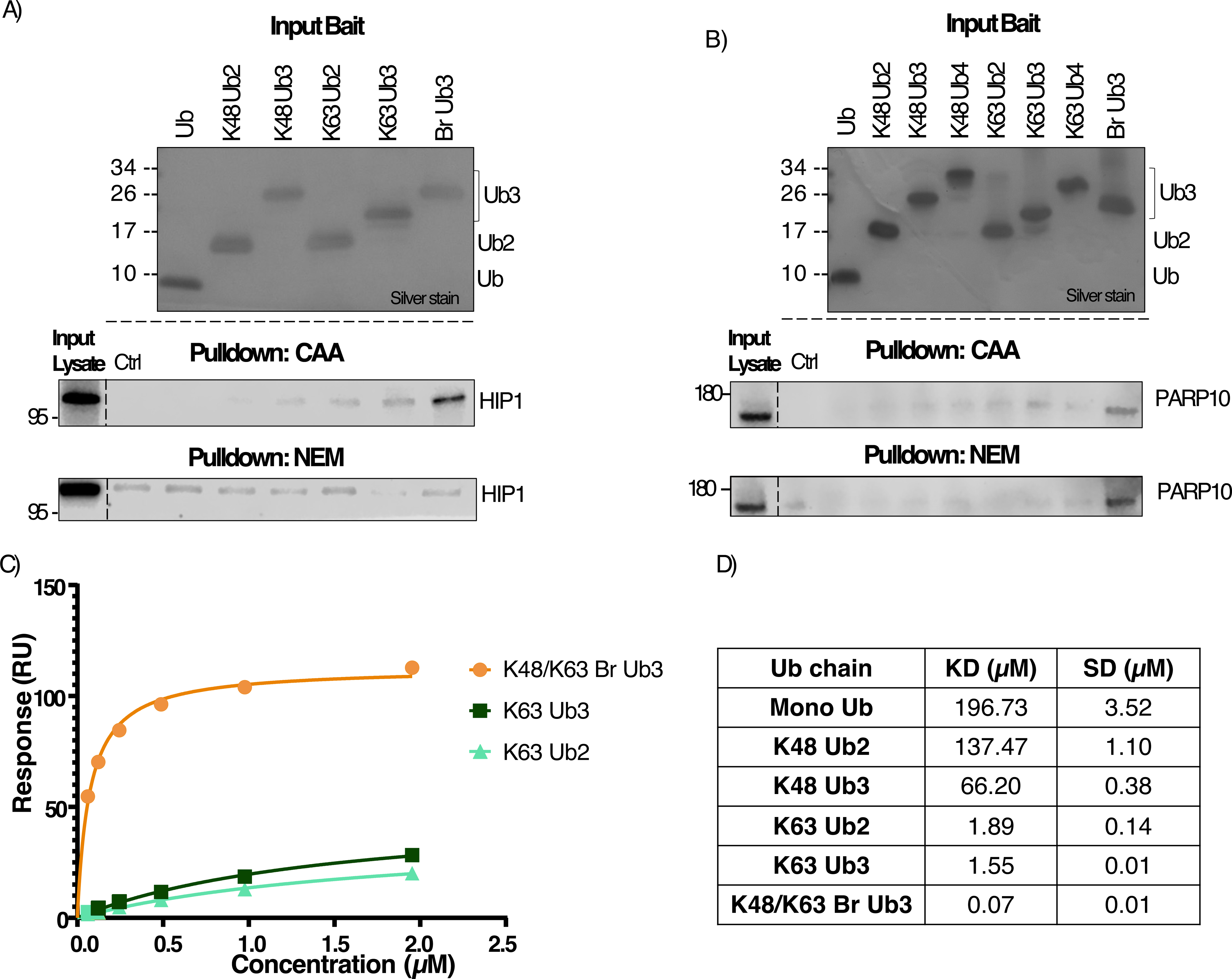
Further validation of Hip1 and Parp10 branch specificity. A) Western blot of HIP1 enrichment on K48/K63-linked branched Ub from CAA and NEM-treated lysate. Same Ub interactor pulldown experiment, as in Figure 1 B, thus same silver stain of input Ub. Blotted with anti-HIP1 antibody. B) Western blot of PARP10 enrichment on K48/K63-linked branched Ub from CAA and NEM-treated lysate. Same Ub interactor experiment as in S.Fig 7 D, thus same silver stain of Ub input. Blotted with anti-PARP10 antibody. C) Overlayed affinity plots of triplicate measurements with determined K_D_ values and respective Standard Deviations (SD) of K63-linked Ub2, K63-linked Ub3 and K48/K63-linked branched Ub3 chains interacting with immobilized biotinylated HIP1 protein. Averaged response values [RU] at equilibrium were plotted against the injected concentration [µM] of respective analytes determined by SPR multi cycle format and fitted according to a steady state affinity model. D) Calculated KD affinities and standard deviations (SD) of Hip1 with Ub chains measured by SPR and fitted via a steady state affinity model.

## Discussion

Our dataset provides a resource of K48- and K63-linked Ub interactors in humans and budding yeast, and their chain length and homotypic versus branched heterotypic binding preference. The linkage-specific interactors we identified included many known K48- and K63-specific UBPs and their associated cellular pathways. Also, amongst the interactors identified with a binding preference for Ub3 over Ub2 were UBPs with multiple Ub-binding sites which are known to bind ≥Ub3. Furthermore, we validated the K48/K63-linked Ub preference of one of our MS hits, HIP1, using SPR binding assays. These results validate our method’s ability to identify chain type-specific interactors.

The length of Ub chains in the cell and the affect this has on Ub-binding is still being elucidated. Interestingly, our data revealed that chain length (Ub3 versus Ub2) has a larger effect on K48-linked Ub-binding, than on K63-linked Ub-binding. This observation could be due to the topologies of different chain linkage types – K48-linked Ub moieties self-associate and have a higher propensity to form more ‘closed’ structures^75–77^, whereas K63-linked Ub chains have a more open ‘beads on a string’ topology^77,78^. Thus, the conformational space of K48-linked chains may be more affected by the addition of Ub moities^79^. Additionally, it was shown in yeast that K63-linkages exist most often in Ub2 chains^80^, which could reduce the need for K63-specific UBPs with preference for longer Ub chains. Whereas K48-linkages were found to exist in both Ub2 and ≥Ub3 chains. We also observed that K63-specific UBPs were more enriched on Ub3 K48-linked Ub chains than K48-linked Ub2. This could be a result of avid binding to non-adjacent Ub moieties^34^ or the wider conformation distribution of >Ub2 K48-linked Ub chains^79^. Interactors enriched on K48-linked Ub3 over Ub2 included UBPs with documented specificity for longer Ub chains: ZFAND2B/AIRAPL^50^, ATXN3^60,61,81^ and MINDY1^41^, which bind 3, 3 and 4 Ub moieties respectively, and DDI2/Ddi1 in yeast^82^. We also identified Ub3 preference for proteins with previously undetermined chain length specificity, CCDC50, APPL1 and FAF2. The molecular mechanisms behind their length preference and the relevance of this for their cellular function is a direction for future study.

Elucidating the interactors of branched Ub chains is a key area of investigation in the Ub field. We observed that linkage-specific Ub interactors also bind K48/K63-linked branched chains. This supports the model of branched chains as a combination of two Ub signals. Contrastingly, it has been suggested that branching can inhibit UBP binding, as reported for K63-specific DUB CYLD^12^. In different statistical comparisons we observed reduced enrichment of known K63-specific UBP TOM1 and K48-specific UBPs RAD23A and RAD23B, across datasets, and DUB USP11, in the CAA dataset, on K48/K63-linked branched Ub. These observations suggest that in some contexts the branch may reduce Ub binding.

The cellular functions of branched Ub interactors can reveal the pathways in which branched Ub chains play a role. Two of the enriched K48/K63-linked Ub interactors identified here, RFC1 and MORC3, were also found in a recent preprint by Lange et al^14^. Interestingly, RFC1 along with PARP10 and USP48, which we also identified as K48/K63-linked Ub specific interactors, are all associated with DNA damage repair^83–85^. In line with this, we previously observed sensitivity to hydroxyurea with a Ubc1 UBA mutant (in a Ubc4 KO background)^19^ and Lange et al observed an increase in K48/K63-linked Ub at the site of DNA damage^14^. Our results also suggest a role of K48/K63-linked branched chains in Huntington’s disease, as we identified the Huntingtin-interacting protein HIP1 and UBR4, which has been linked to K11/K48-linked branched Ub at mutant Huntingtin^66^, as branch-specific interactors. Finally, two of our K48/K63-linked Ub specific interactors are E3 ligases, UBR4^13,66^ and MIB2^70^ and one is a Cullin RING ligase substrate-adaptor, ANKFY1^69^. The branchpoint may, therefore, be a scaffold for further ubiquitination.

The Ub chain type-specificity of UBPs in yeast is less well characterised. We provide the first resource of chain-specific Ub interactors in budding yeast. Our data is supported by the identification of known chain linkage-specific UBPs. We also identify the linkage-type specificity of expected UBPs with previously unknown chain preference, including K48-binding Cia1 and K63-binding Duf1 and Gts1. We reproduced the finding of Ddi1 length preference and showed that this specificity was conserved from yeast to humans.

As our screen utilised native Ub chains, it was essential that we blocked Ub chain disassembly by inhibiting endogenous DUBs. We provide the first comparison of cysteine alkylators NEM and CAA as DUB inhibitors. We observed that lysate treatment with NEM stabilised Ub chains, better than treatment with CAA, in line with literature on the potency of NEM^21^. However, potentially due to its unspecific alkylation activity^25^, NEM also inhibited the Ub binding of some UBPs, including HIP1, or altered their chain-type specificity, as for IKBKG/NEMO. These findings bring caution to the use of unspecific alkylating reagents in protein binding studies. For DUB inhibition more specific reagents are available; inhibitors against specific DUBs^86^ or for broad-range inhibition of DUBs PR-619^87^, however it is weaker against non-cysteine protease DUBs^88^.

In summary, this resource provides readers, interested in the Ub code, with the chain type, linkage, length and homotypic versus heterotypic, preference of K48- and K63-linked Ub interactors in humans and yeast. We were able to identify known linkage- and length-specific UBPs, enhancing the reliability of our method. The significant chain-type enriched proteins we identified serve to inform potential models of Ub signalling pathways on which further experiments can be based.

## Materials and Methods

### Plasmids and reagents

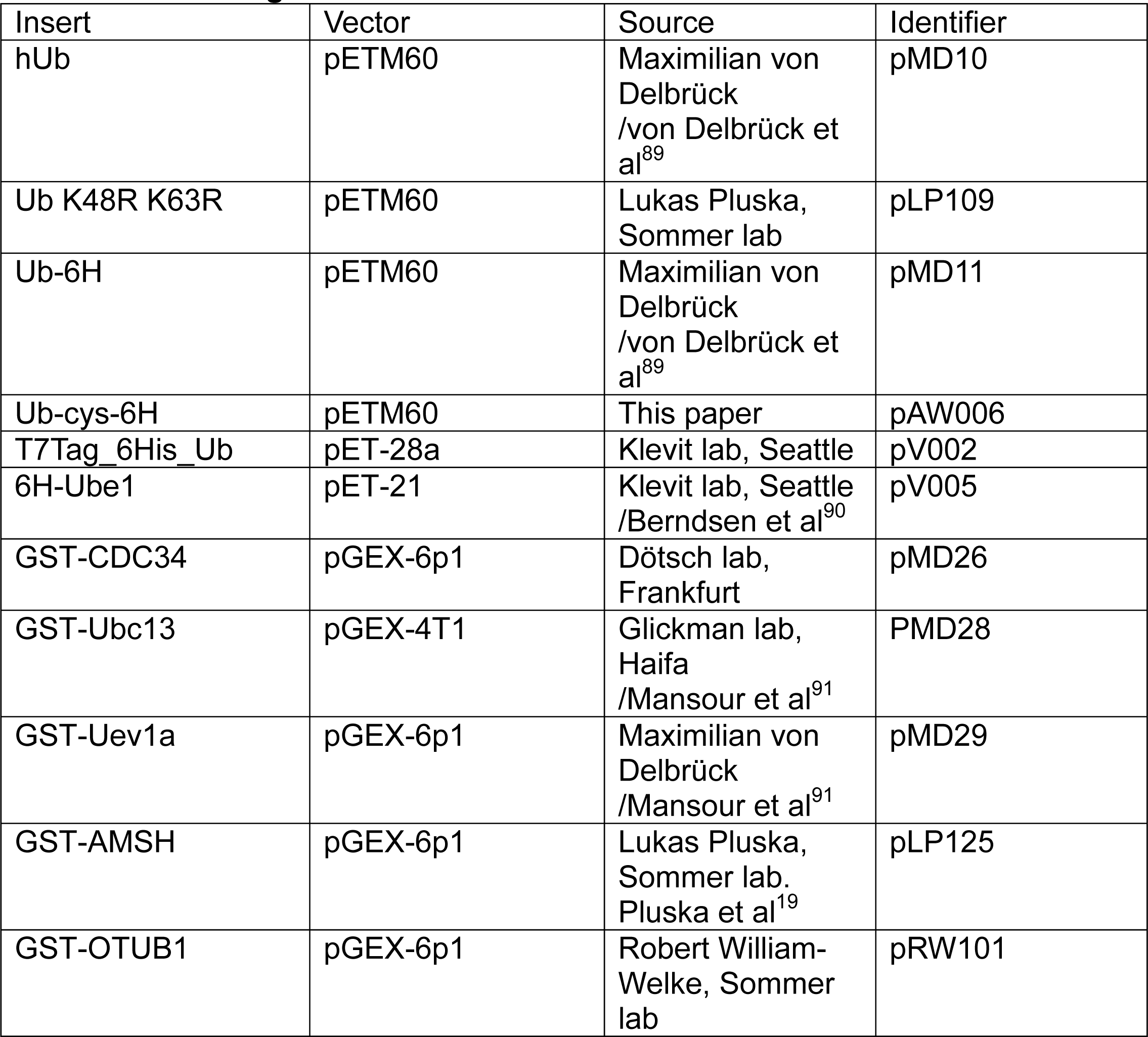

### Cloning

A glycine serine repeat linker containing one cysteine residue was cloned between the ubiquitin (Ub) C-terminus and 6 x histidine tag in pMD11 (hUb-6H in pETM60) using Gibson Assembly.

### Recombinant protein purification

BL21 Rosettta cells were transformed with expression plasmids and grown overnight at 37°C. At an OD600 of 0.8-1.0 the culture was cooled to 18C and expression was induced with 0.5mM IPTG and incubated overnight. Cells were pelleted and resuspended in lysis buffer: 50mM TRIS pH 7.5 150mM NaCl 1mM PMSF. Cells were then lysed with an Avestin EmulsiFlex-C5 homogenizer and cleared of cell debris by centrifugation at 20,000 g at 4°C for 20 minutes.

For purification of 6xhis-tagged Ubs, 5mM Imidazole and 3mM 2-mercaptoethanol were included in the lysis buffer. Lysate was incubated with TALON metal affinity resin (Takara) (3ml slurry per L culture) for 1 hour at 4C on a rotor. The resin was washed in a gravity column (BIORAD) with 4 x wash buffer: 50mM TRIS pH 7.5 150mM NaCl 3mM 2-mercaptoethanol 10mM Imidazole. Protein was eluted with elution buffer: 50mM TRIS pH 7.5 150mM NaCl 300mM Imidazole. A final concentration of 2mM DTT was added.

For purification of GST-tagged proteins, 4mM DTT was added to cleared lysate, which was then incubated with Glutathione Sepharose 4B beads (Cytiva – formerly GE Healthcare) (5ml per L culture) for 1 hour at 4°C. The resin was washed in a gravity column with wash buffer: 4 x 50mM TRIS pH 7.5 150mM NaCl 4mM DTT. Protein was then eluted with elution buffer: 50mM TRIS pH 7.5 150mM NaCl 20mM Glutathione.

Purification of untagged Ub was performed by acidic preparation^92^. 70% perchloric acid was added dropwise under a fume hood to cleared lysate until the solution becomes very cloudy. The precipitant was removed by centrifugation at 20,000 g at 4C for 20 minutes. The supernatant was neutralised to pH 7.5 using 10M NaOH.

All proteins were further purified by size-exclusion chromatography. Protein solutions were concentrated using Amicon Ultra centrifugal filter (Millipore) for application to FPLC AKTA pure (GE Heathcare). For Ub purification a HiLoad 26/600 Superdex 75 pg column (Cytiva) was used. For larger protein Superdex 75 increase 10/300 GL column (Cytiva) was used. Proteins were checked by SDS-PAGE and Coomassie staining. Protein concentration was determined using a DC protein assay (Biorad).

### Active UBE1 purification

A Ub-affinity-gel column was prepared in order to purify active bound UBE1. All steps carried out at 4C, unless stated otherwise. 5ml Affi-Gel 19 slurry (BIORAD) was added to a gravity column (BIORAD) and washed 3 x 10ml H2O. 6ml of 50mg/ml 6His-T7tag-Ub purified in MOPS buffer was added and incubated overnight at 4C on a roller. The supernatant was removed, then the unconjugated resin was blocked by incubation with 0.5ml 1M ethanolamine pH 8 for 1 hour on roller. The column was washed with 6 x 0.1M 3-(N-morpholino)propanesulfonic acid (MOPS) pH 7.2 and stored at 4C in 0.1M MOPs 2% sodium azide pH 7.2 until use.

UBE1 was expressed as in 4.2.2.1 and then lysed in 50mM TRIS pH 8 0.2mM DTT 10mM MgCl_2_ 200uM PMSF with an Avestin EmulsiFlex-C5 homogenizer. Lysate was cleared of cell debris by centrifugation at 20,000 g at 4°C for 20 minutes and then further filtered through 0.45micron filter. Storage buffer was removed from the Ub-affinity-gel column and it was equilibrated in 50mM TRIS HCl pH 8. Filtered UBE1 lysate was added to the prepared Ub-affinity-gel column. The column was topped up with 5ml ATP and MgCl_2_ pH 7 to final concentrations of 40mM of each. The column mix was incubated at RT for 1 hour to conjugate the UBE1. The flowthrough was removed and the column was washed approximately 20 x with 50mM TRIS-HCl pH 8 0.5M KCl (until no contaminants at 280nm by Nanodrop). 5ml elution buffer 50mM TRIS HCl pH 8 10mM DTT was added and incubated for 10 minutes before collecting flowthrough. This step was repeated 6 times.

Pooled eluate was dialysed into 50mM TRIS HCl pH 8 150mM NaCl 1mM DTT overnight, using two dialysis steps with sealed dialysis tubing and 4L buffer. The final sample was concentrated with an Amicon Ultra-15 filter (MWCO 50).

The Ub-Affinity-gel was washed and stored in 0.1M MOPs with 2% sodium azide. It was regenerated for further use by washing 3 x 50mM TRIS HCl pH 9 1M KCl.

### Preparation of Ub chains

Ub chains in which the proximal Ub contains a C-terminal cysteine followed by a 6xHis tag were assembled *in vitro* using recombinant Ub ligases. For homotypic chain synthesis, the reaction mix contained E1 ligase 2μM hUbe1, E2 ligases 25μM Cdc34 or 6μM Uev1a and 6μM Ubc13 for K48-linked or K63-linked chain synthesis, respectively, 1.2mM Ub and 0.8mM 6xHis-cys-Ub. K48/K63-linked branched chains were synthesised with 2μm hUbe1, 10μM Ubc1, 8μM Uev1a, 8μM Ubc13, 0.5mM 6xHis-cys-Ub and 1mM K48R, K63R Ub. Synthesis reactions were performed in 50mM Tris-HCl pH 8 9mM MgCl2 15mM ATP 5mM Beta-mercaptoethanol with a total volume of 3-5ml. Reactions were incubated overnight at 37°C. The reaction was diluted with 50mM TRIS pH 7.5 150mM NaCl 7.5mM Imidazole 1.5mM BME. 6xHis-tagged chains were then purified with TALON resin (5ml per 1ml reaction) and washed 3 x wash buffer: 50mM TRIS 150mM NaCL 3.5mM 2-mercaptoethanol 5mM Imidazole pH 7.5. After elution with 300mM Imidazole, Ub chains of different lengths were separated by size exclusion on a to FPLC AKTA pure (GE Heathcare) using a HiLoad 26/600 Superdex 75pg column (Cytiva), with a low flow rate (0.3-0.5 ml/min). For future immobilisation of Ub chains, biotin was conjugated onto the proximal Ub of the chain via maleimide-cysteine reaction. 0.5-1.5mg Ub chain was reduced by incubation for 1 hour at 37C with 10 x molar excess of tris(2-carboxyethyl)phosphine (TCEP). Reducing agents were then removed by filtering through Pierce dye and biotin removal column (Thermo Scientific). The resulting chains were then incubated overnight at room temperature with 10x molar excess of EZ-Link Maleimid-PEG2-Biotin (Thermo Scientific). The next day, excess biotin was quenched with a final concentration of 10mM DTT. Excess biotin and DTT was removed by 6 x sequential dilution and concentration using 3KDa Amicon Ultra centrifugal filter (Millipore). Yield of biotin conjugation was tested by intact MS with an Agilent 6230 b LC/MS TOF mass spectrometer. Chain concentration was determined by DC protein assay (Biorad).

### Cell culture / Lysate preparation

For Ub interactor copulldown, wildtype *S. cerevisiae,* yeast and HeLa lysate were prepared.

5L yeast culture was grown to 1 OD600, harvested at 4,000 x g for 5 minutes, washed in 25ml water 1mM PMSF and frozen -80°C until further use. Cell pellet was resuspended in yeast lysis buffer (200μl per 100ml culture): 50mM TRIS-HCl pH 8 150mM NaCl 0.4% NP40 (IPEGAL ca630) 1mM PMSF 5% glycerol and aliquoted into 2ml Eppindorf tubes for lysis. Glass beads (Carl Roth) (≈ 200μl per tube) were added and cells were lysed on with a vortex at max. speed for 5 x 1 minute (incubation on ice in between vortexing). Minimal lysis buffer: 50mM TRIS pH 8, 150mM NaCl 0.1% NP40 10mM CAA 1mM PMSF (300μl per tube) was added. Cell debris was removed by centrifugation at 3000 rpm for 3 minutes at 4°C. Supernatant was transferred to a fresh 1.5ml Epindorf tube and lysate was cleared by centrifugation at 20,000 x g for 5 minutes at 4°C. Pierce BCA assay (Thermo Scientific) was used to measure protein concentration of the lysate.

HeLa cells were grown in Dulbecco’s modified Eagle’s medium (DMEM), supplemented with 10% fetal bovine serum (FBS) and 1% Penicillin/Streptomycin at 37°C, 5% CO_2,_ and 90% humidity. 50 x 15cm dishes HeLa cells were harvested with Trypsin (3ml per 15cm dish) at 90% confluency. Cell pellets were washed with 4 x Phosphate-buffered Saline (PBS) (Sigma Aldrich) and cell pellets were frozen at - 80C for further use. Cells were resuspended in Hela lysis buffer (450μl per 15cm dish): 150 mM NaCl, 50mM Tris-HCl, pH 8.0, 0.5% (vol/vol) IGEPAL, 5% (vol/vol) glycerol 1mM PMSF 1x protease inhibitors mix, briefly vortexed and incubated for 45 minutes at 4C on a rotor. Lysate was centrifuged at 14,000 x g for 15 minutes at 4°C. Pierce BCA assay (Thermo Scientific) was used to measure protein concentration of the lysate. Before use in Ub interactor pulldown, 1mM N-Ethylmaleimide (NEM) or 10mM Chloroacetamide (CAA) was added as a deubiquitinase inhibitor and incubated with lysate for 1 hour.

### Ub interactor pulldown for mass spectrometry

Ub interactor pulldown was done in quadruplicate for each chain type or resin only control. 25-50μg Ub chains were immobilised on streptavidin magnetic sepharose resin (Cytiva) by incubation on rotor for 1 hour at 4°C in 200μl binding buffer (50 mM Tris pH 8.0, 150 mM NaCl, 0.1% NP40). Immobilised chains were washed once with binding buffer. 2.5-4mg Hela or yeast lysate was added to immobilised chains and incubated overnight on rotor at 4°C. Next day, resin was washed 3 x with 1ml wash buffer (50mM TRIS pH 8, 150mM NaCl). Enriched material was eluted by on bead digest, as detailed below.

### Sample preparation and mass spectrometry

Sample preparation was conducted using on-bead tryptic digestion, adhering to the protocol established^93^. Briefly, washed beads were incubated in digestion buffer: 2 M urea, 50 mM Tris (pH7), 1 mM dithiothreitol (DTT) and 0.4 µg sequencing grade trypsin (Promega) for 1 h at 25°C with continuous agitation on a shaker operating at 1000 rpm. Following the incubation period, the supernatant was carefully transferred to a fresh tube. The beads were washed again 2 x with urea/Tris buffer and each time combined with the supernatant with previous steps. Proteins were reduced with 4 mM DTT for 30 min and alkylated using 10 mM CAA for 45 min at 25°C whilst shaking at 1000 rpm. Proteins were subjected to a second digest overnight with 0.5 µg trypsin and incubated at 25°C whilst shaking at 700 rpm. Following the tryptic digestion, we employed stage-tips, following the procedure described^94^ to remove salts and impurities from the samples. For human CAA-treated samples, peptides were cleaned-up using a peptide-based SP3 approach^95^. Briefly, peptides in acqueous solution were incubated with SP3 beads at ratio of 200:1 beads:protein. ACN was added to a final concentration of 95% and beads were were subsequently washed 3 x with 100% ACN. Peptides were eluted from bead in 50 µl LC-MS grade water.

The samples were then subjected to liquid chromatography-mass spectrometry (LC-MS) measurements using an orbitrap Exploris 480 mass spectrometer (Thermo Fisher Scientific) in conjunction with an EASY-nLC 1200 system (Thermo Fisher Scientific). The mass spectrometer was operated in data-dependent mode, and a 110-minute gradient was applied.

### Proteomic Analysis

For the analysis of mass spectrometry data, we utilised MaxQuant version 2.0.3.0^96^ incorporating MaxLFQ-based quantitation^97^ and enabling the match-between runs algorithm. Carbamidomethylation of cysteine residues, acetylated protein N-termini and oxidised methionine were designated as variable modifications. Peptides containing cysteine residues not used in quantitation. The Andromeda search was performed using a Uniprot human or yeast database from 2022 including protein isoforms, along with a list of common contaminants. An FDR cutoff of 0.01 was applied on the PSM and protein level.

Subsequent to data acquisition and initial processing, downstream data analysis was conducted in the R programming environment (v4.2.1) using iBAQ values for quantitation In R (v4.2.1), data was reverse filtered, dropout samples were removed and IBAQ values were used to filter by valid values (≥3 per chain type), median normalised and imputed using a normal distribution with downshift approach. For use in further analysis, Ub enriched proteins were pre-filtered by significance in at least one Ub pulldown in a two moderated T-test against the control bead-only pulldown (adjusted P value < 0.05, log (fold change) < 0). This filter was then applied to data at the step before normalisation and imputation. Subsequently filtered data was median normalised and imputed using a normal distribution with downshift. Data that has undergone this prefiltering is described as pre-filtered Ub enriched proteins in text and figure legends.

Statistical tests were performed in R using the Shiny app ProTIGY provided by the Broad Institute on GitHub (https://github.com/broadinstitute/protigy). Moderated F test was used to compare all pulldown samples. Two sample moderated T-test was used for pairwise comparisons. To identify significant hits a significance a threshold of < 0.05 was applied for Benjamini Hochberg corrected adjusted P values (adj.P), unless otherwise stated. Data visualisation was performed in R using the tidyr, dplyr, tibble, tidyverse, pheatmap, ggplot2, stringr, corrplot and VennDiagram packages. Metascape was used for gene ontology (G0) enrichment ^98^ with express analysis settings: minimum overlap of 2, P Value < 0.01 and minimum enrichment of 1.5.

### Ub interactor pulldown validation for Western Blot

14μg Ub chains were immobilised on streptavidin magnetic sepharose resin (Cytiva) by incubation on rotor for 1 hour at 4°C in 100μl binding buffer (50 mM Tris pH 8.0, 150 mM NaCl, 0.1% NP40). Immobilised chains were washed once with binding buffer. 1mg Hela or yeast lysate was added to immobilised chains and incubated overnight on rotor at 4°C. Next day, resin was washed with 3 x 0.5ml wash buffer (50mM TRIS pH 8, 150mM NaCl). Pulled-down material was eluted in 25μl SDS sample buffer with 2-merpcatoethanol and boiled at 95C for 10 minutes. 10μl samples were run on SDS-PAGE and Western Blot against Ub and proteins of interest.

### Western Blot

SDS-PAGE was done using homemade Bis-acrylamide gels or mini protean TGX gradient gels (BioRad). Gels were blotted onto PVDF membranes. Membranes were blocked in 5% milk powder or 10% roti block (Roth). Primary antibodies were added overnight at 4C in 5% milk powder or 10% roti block (Roth) – 1:1000 Ub antibody P4D4 (Santa Cruz Biotechnology), 1:3000 Hip1 (Proteintech 22231-1AP), 1:000 PARP10/ARD10 (Novus Biologicals NB100-2157), 1:1000 Rad23b (Elabscience E-AB-62188), 1:1000 Dsk2 (Abcam ab4119-100), 1:4000 Rad23 (Sommer lab), 1:1000 Epn2 (Invitrogen), DDI2 (Abcam AB197081), 1:1000 DDI1 serum (Jeffery Gerst, Weizmann Institute of Science^99^), 1:1000 APPL1 (Cell signalling D83H4), 1:1000 CCDC50 (Abcam AB127169), 1:1000 FAF1 (Proteintech 1027-1-AP), 1:1000 ZFAND2B (Sigma Aldrich), 1:1000 Riok3 (Proteintech 13593-1-AP), 1:1000 USP11 (Proteintech 22340-1-AP), 1:5000 CDC48 (Sommer lab), 1:1000 Vps9 (Scott Emr, Cornell University^100^), 1:1000 YUH1 serum (Tingting Yao, Colorado State^101^). Anti-mouse IgG HRP (Sigma) and Anti-rabbit IgG HRP (Sigma) were used 1:10,000 as secondary antibodies. Immunoblots were visualized using an Odyssey XF Imager (Li-Cor).

### Ub chain disassembly (UbiCREST)

1.25ug ub chains were incubated with 1uM deubiquitinase, either K48-specific OTUB1 or K63-specific AMSH, in disassembly buffer (50 mM Tris-HCl pH 7.5, 50 mM NaCl, 10 mM DTT) for 45 minutes at 37C. Reactions were stopped with SDS DTT sample buffer. Samples were run on SDS-PAGE, Western Blot with anti-Ub antibody and imaged using an Odyssey XF Imager (Li-Cor).

### Surface Plasma Resonance

SPR experiments were performed on a Biacore T200 at 25°C using 50 mM HEPES pH 7.5, 300 mM NaCl, 0.5 mM TCEP, 0.05% Tween 20 and 1% BSA as the running buffer. Series S Sensor Chips NA (Cytiva) were used according to the manufacturer’s recommendations: 3x injection of 50 mM NaOH, 1 M NaCl for 60 s to condition the surfaces with a subsequent injection of recombinant biotinylated HIP1 protein at 2 µg/mL diluted in running buffer. A final response of 900 RU was reached for three flow channels by immobilizing the protein for 90 s at 10 µl/min. One flow channel was used as an empty reference surface without protein injection.

Each Ubiquitin (Ub) analyte was measured at a flow rate of 30 µl/min using multi cycle conditions injecting single concentrations during each cycle increasing from 60 nM up to 250 µM over the reference and protein surfaces. Resulting Sensorgrams were referenced, blank subtracted and evaluated.

Evaluation was carried out by using the steady state affinity model:

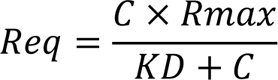

where C is the injected concentration, Rmax is the maximal response and Req is the response at steady state to determine respective K_D_ values.

## Acknowledgements

We would like to thank some of our colleagues at the Max Delbrück Centre for Molecular Medicine: Ernst Jarosh for his advice during the project and manuscript proofreading, Mandy Gerlach for her technical assistance with Western Blotting, Robert William Welke for providing OTUB1 and AMSH DUBs, Sabine Meyer and Anja Schutz from the protein production facility for their technical assistance with the LC/MS TOF mass spectrometer and Mohamed Haji from the proteomics facility for his technical assistance with the LC/MS mass spectrometer. We would also like to thank Mahil Lambert at the University of Frankfurt for his assistance in protein purification. Additionally, thanks to those who kindly gifted us antibodies.

We would like to acknowledgment our funding agencies. The joint DFG grant DO 545/17-1 | SO 271/9-1. Also, the Structural Genomics Consortium (SGC), a registered charity (No:1097737) that received funds from Bayer AG, Boehringer Ingelheim, Bristol Myers Squibb, Genentech, Genome Canada through Ontario Genomics Institute, Janssen, Merck KGaA, Pfizer, and Takeda. This project received funding from the Innovative Medicines Initiative 2 Joint Undertaking (JU) under grant agreement No. 875510. The JU receives support from the European Union’s Horizon 2020 research and innovation programme, EFPIA, Ontario Institute for Cancer Research, Royal Institution for the Advancement of Learning McGill, University, Kungliga Tekniska Hoegskolan, and Diamond Light Source Limited. Disclaimer: This communication reflects the views of the authors, and JU is not liable for any use that may be made of the information contained herein.

## Author Contribution

L.P and A.W conceptualised the study. A.W cloned plasmids, prepared proteins and Ub chains, maintained cell cultures and performed Ub interactor enrichment pulldowns under supervision of L.P and T.S. O.P performed mass spectrometry under supervision of P.M. A.W and O.P performed downstream analysis of mass spectrometry data. C.L performed SPR experiments and determined binding affinities under supervision of S.K and V.D. A.W wrote the manuscript with input from other authors.

## Figure legends

**Supplementary Figure 1:**
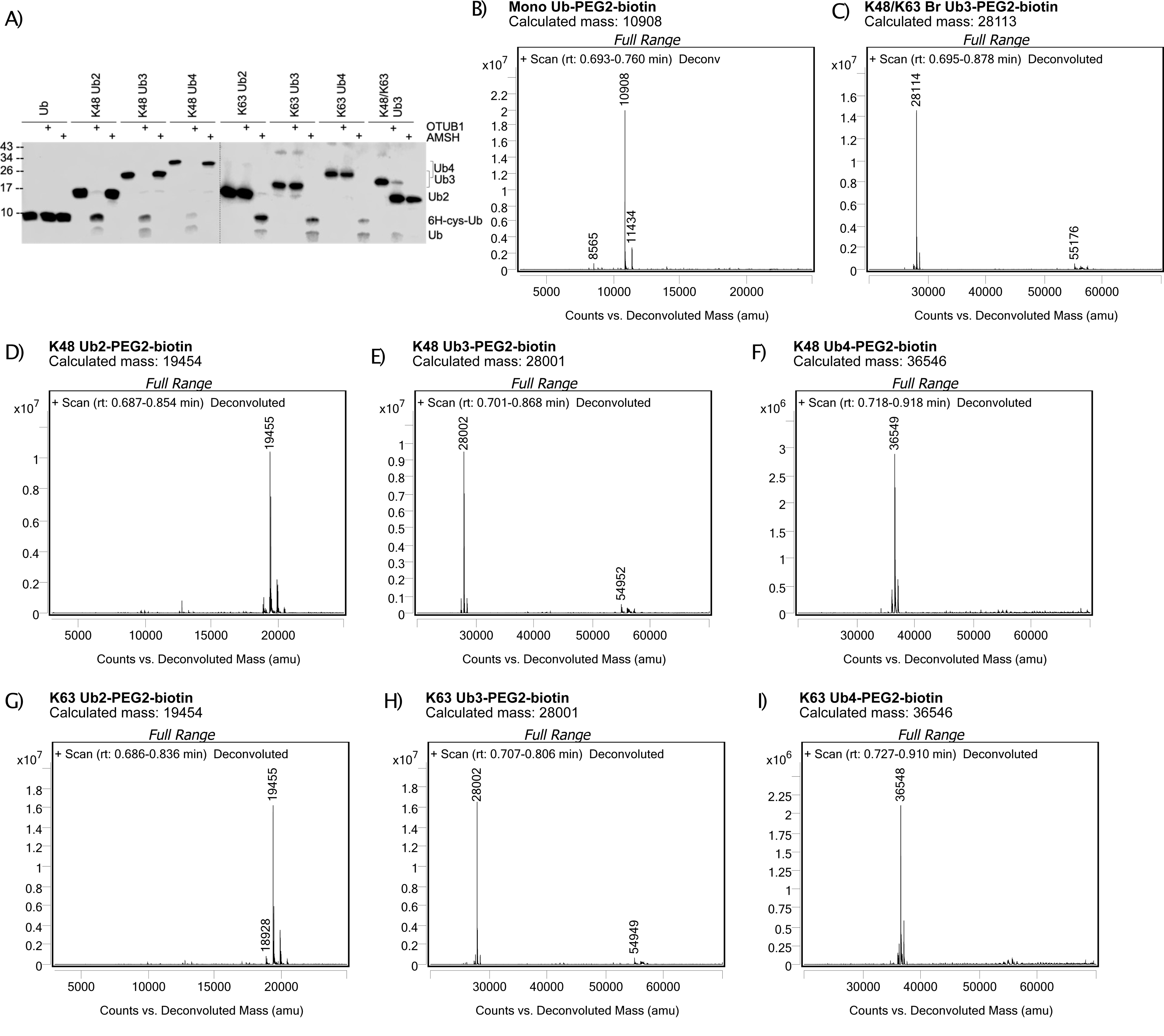
Ub chain quality control. A) Western blot of cleavage of Ub chains by K48- and K63-specific deubiquitinases, OTUB1 and AMSH, respectively. Blotted with anti-Ub antibody. B-I) Intact mass spectrometry deconvoluted spectra of (B) mono Ub, (C) K48/K63-linked branched Ub3, (D) K48-linked Ub2, (E) K48-linked Ub3, (F) K48-linked Ub4, (G) K63-linked Ub2, (H) K63-linked Ub3 and (I) K63-linked Ub4 after Biotin-PEG2-maleimide conjugation.

**Supplementary Figure 2:**
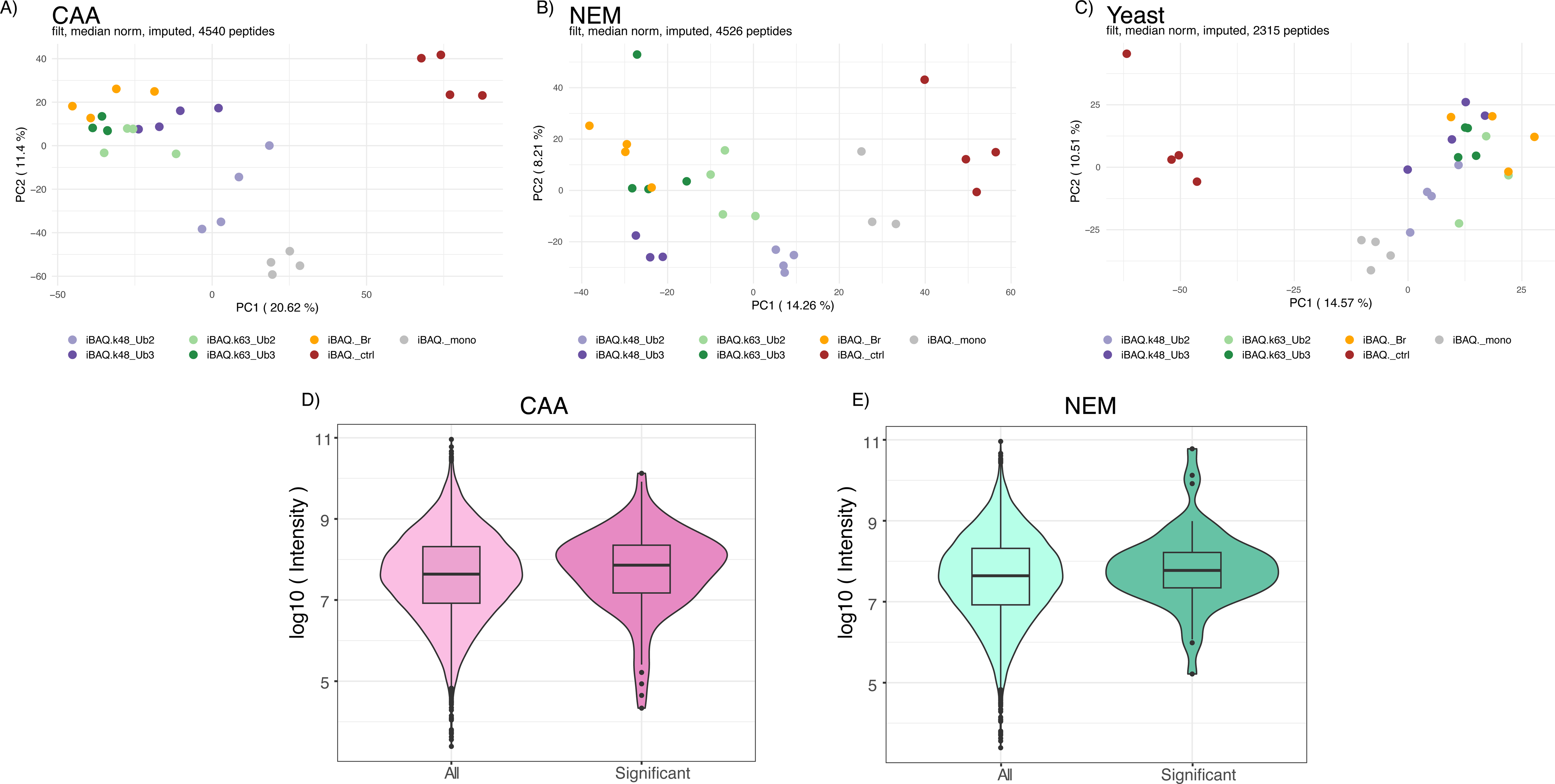
Ubiquitin interactor enrichment-MS quality control. A-C) Principal Component Analysis (PCA) of ubiquitin interactor pulldown samples from (A) CAA-, (B) NEM-treated HeLa lysate and (C) CAA-treated yeast lysate. Each axis in the plot represents a principal component, with percentage of variance explained by each component indicated. D) Protein abundance in whole proteome of all proteins compared to significant hits from ubiquitin interactor pulldown. Intensity values after tryptic digest from Nagarjuna et al deep HeLa proteome used for protein abundance. Isoforms and proteins not identified in deep proteome not included. Significant differently enriched proteins from CAA- and NEM-treated lysate (as in Fig 2A, moderated F test, adj.P cut off < 0.05).

**Supplementary Figure 3:**
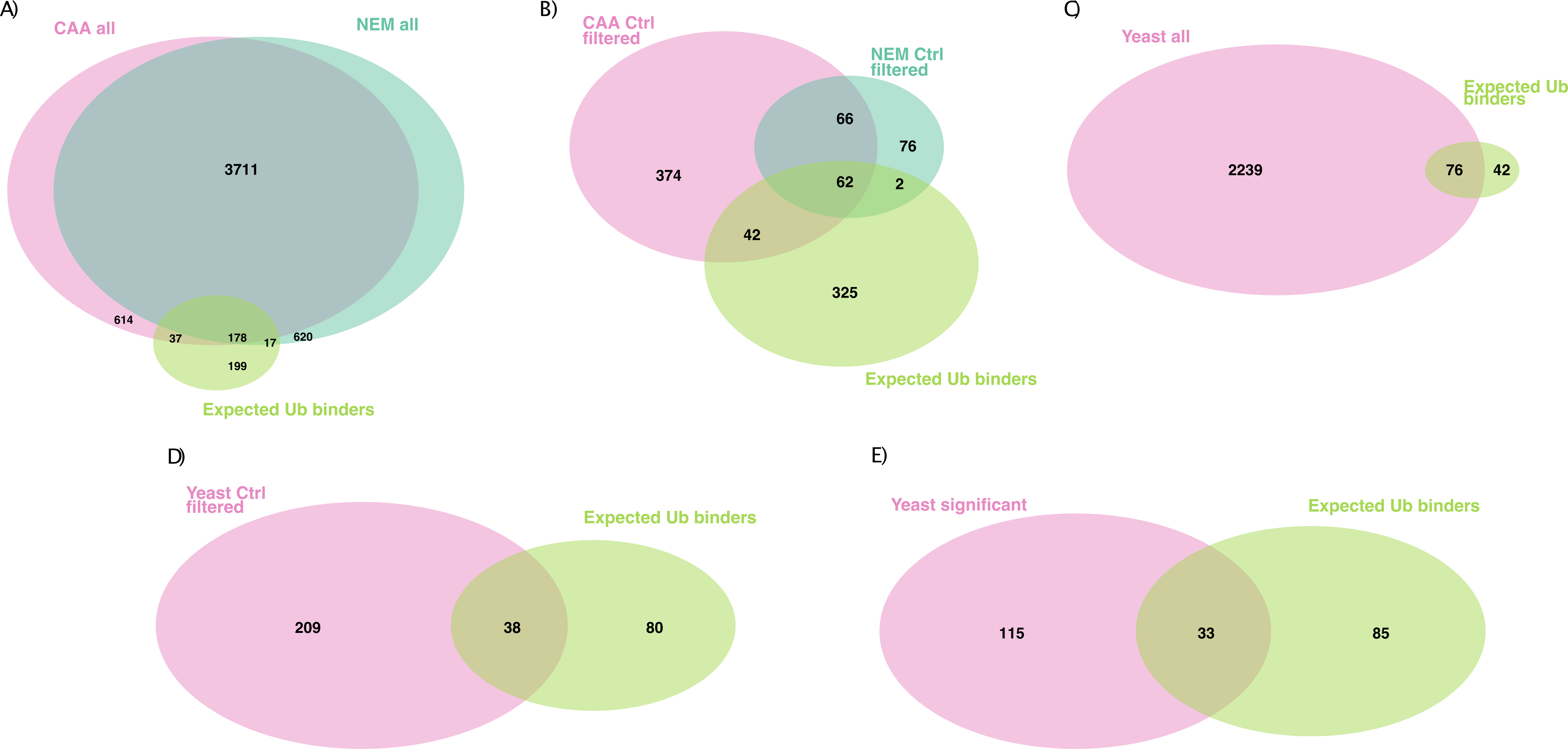
Expected UBPs identified. A and B) Overlap of proteins identified from CAA- or NEM-treated HeLa lysate with expected binders (UBPs). C-D) Overlap of proteins identified from yeast lysate with expected UBPs. (A and C) All proteins identified by MS, (B and D) prefiltered proteins significantly enriched on Ub over the bead-only control (moderated T test, LogFC > 0, Adj.P < 0.05) or (E) significant differently enriched proteins after prefiltering (moderated F test, Adj.P < 0.05). Expected binder lists complied from Gene Ontology term Ub-binding 0043130 and UBD-containing proteins from the UUICD database.

**Supplementary Figure 4:**
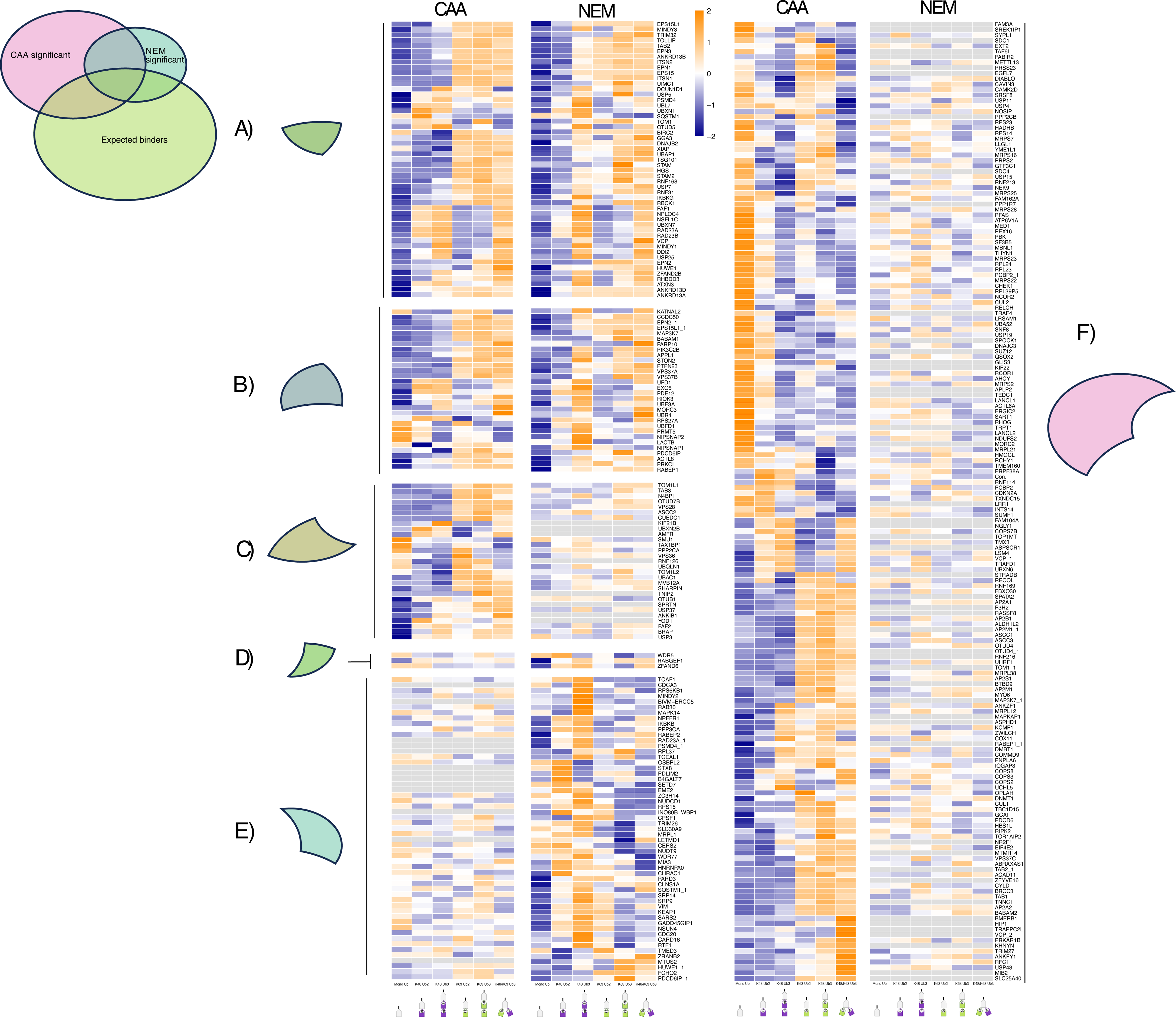
Enrichment pattern of significant proteins across datasets. A-F) Heatmap of enrichment pattern of (A) expected Ub binders which are statistically significant in both datasets, (B) proteins that are significant in both datasets, but not expected ub binders, (C) expected Ub binders that are only significant in the CAA dataset, (D) expected Ub binders that are only significant in the NEM dataset, (E) proteins that are only significant in the NEM dataset and not expected Ub binders and (F) proteins that are only significant in the CAA dataset and not expected Ub binders. Significance is calculated by moderated F test after Ub enriched prefiltering (as in Fig2A, Adj.P cut off < 0.05). Heatmap made using z-scored, moderated F values from moderated F test of all samples, excluding control, before prefiltering. Low transparency of heatmap used to indicate when Adj.P > 0.05, therefore insignificant. Proteins hierarchical clustered by Euclidean distance in the CAA dataset and then ordered the same in NEM dataset, or vice versa when proteins are significant in NEM dataset only.

**Supplementary Figure 5:**
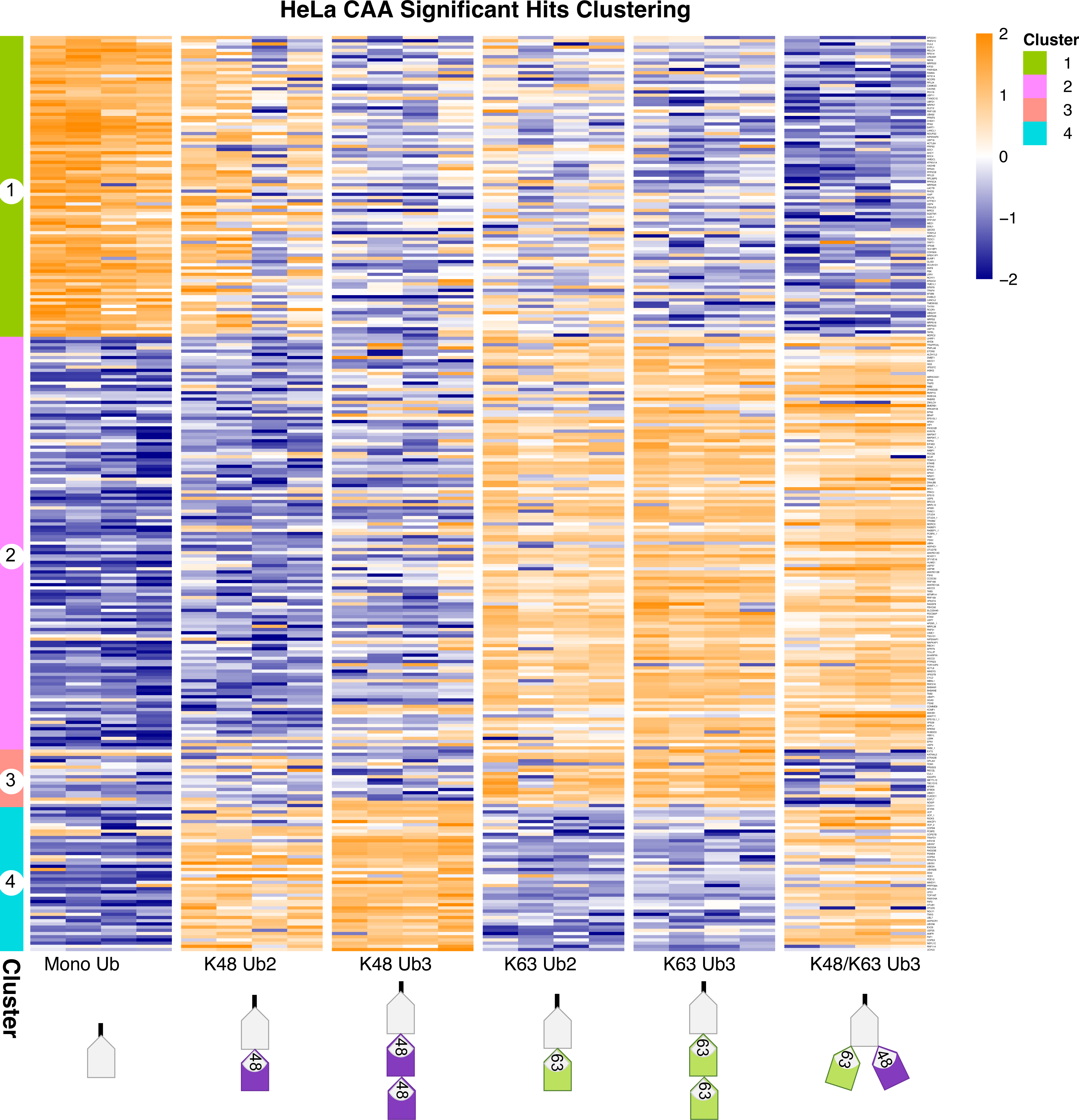
Clustering of significant hits in CAA dataset. Clustered heatmap of significant hits in CAA dataset, identified by moderated F test after Ub enrichment prefiltering (Adj.P cut off < 0.05). Hierarchical clustering by Euclidean distance. Same as in Figure 2C, but with protein names on rows.

**Supplementary Figure 6:**
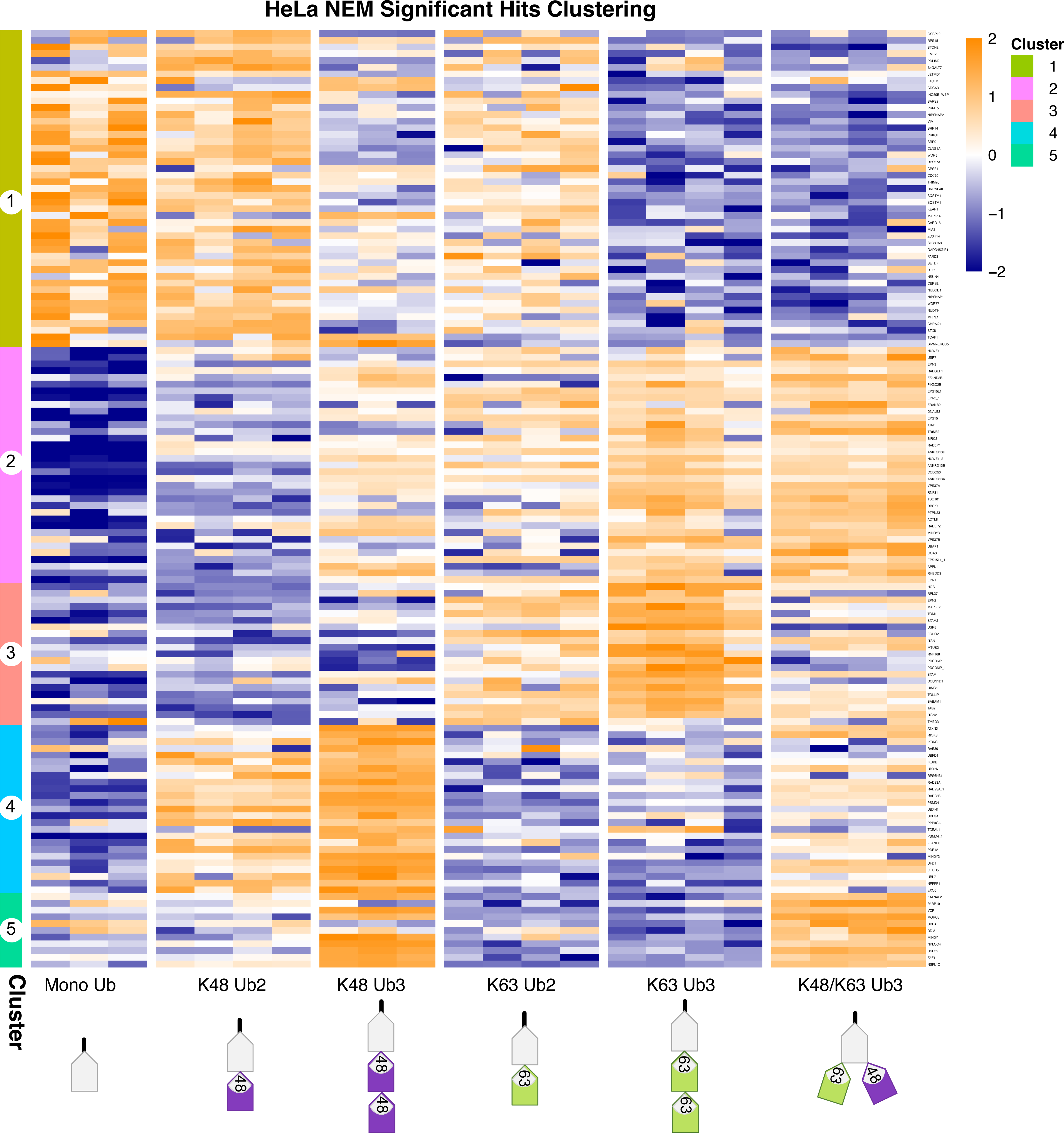
Clustering of significant hits in NEM dataset. Clustered heatmap of significant hits in NEM dataset, identified by moderated F test after Ub enrichment prefiltering (Adj.P cut off < 0.05). Hierarchical clustering by Euclidean distance. Same as in Figure 2D, but with protein names on rows.

**Supplementary Figure 7:**
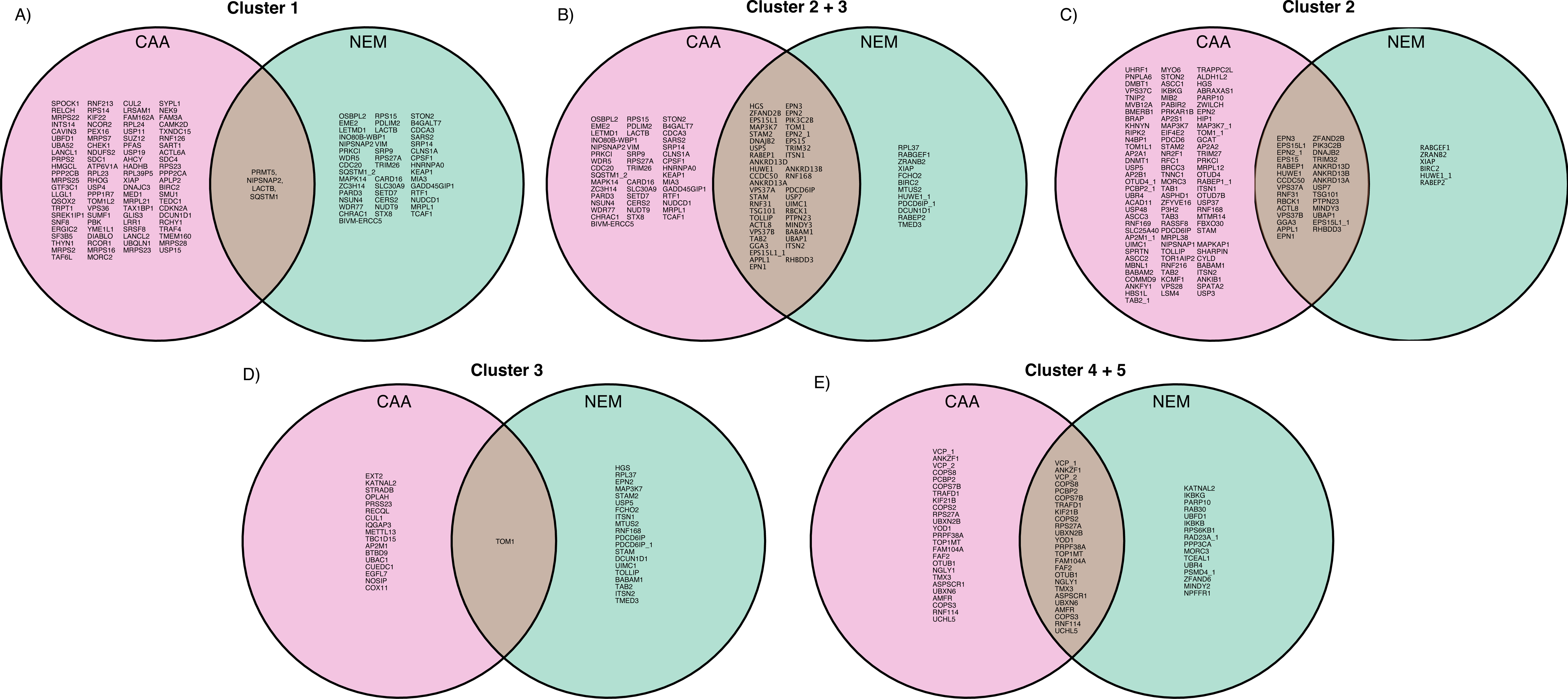
Chain linkage-enriched cluster overlaps between datasets. A, B, C, D and E) Venn diagrams of proteins from (A) Cluster 1, (B) Cluster 2, (C) Cluster 3, (D) Cluster 2 and 3 combined and (E) Cluster 4 and 5 combined between CAA and NEM datasets. Clusters from Figures 2C and D and Supplementary Figures 5 and 6.

**Supplementary Figure 8:**
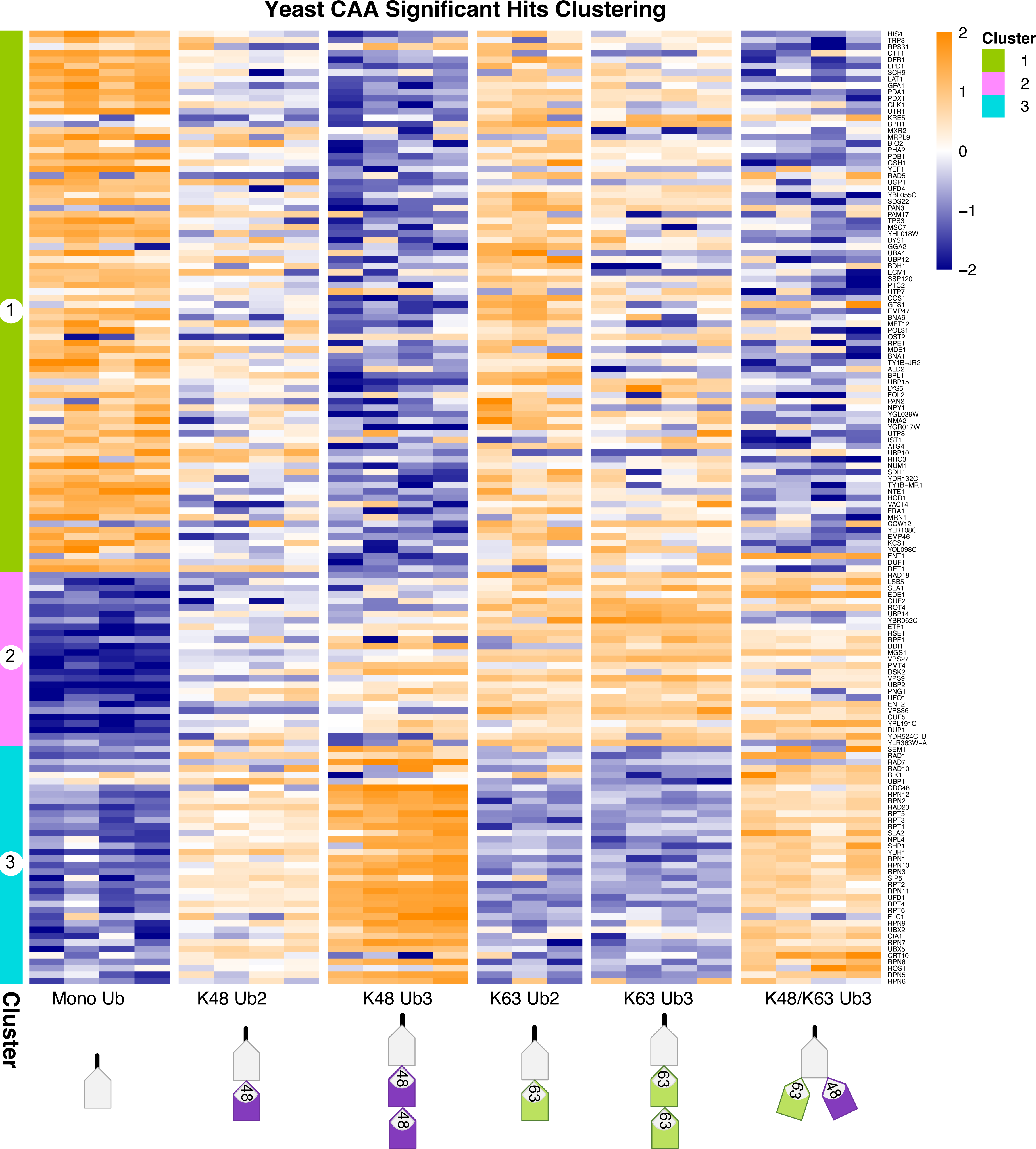
Clustering of significant hits in yeast dataset. Clustered heatmap of significant hits in yeast dataset, identified by moderated F test after Ub enrichment prefiltering (Adj.P cut off < 0.05). Hierarchical clustering by Euclidean distance.

**Supplementary Figure 9:**
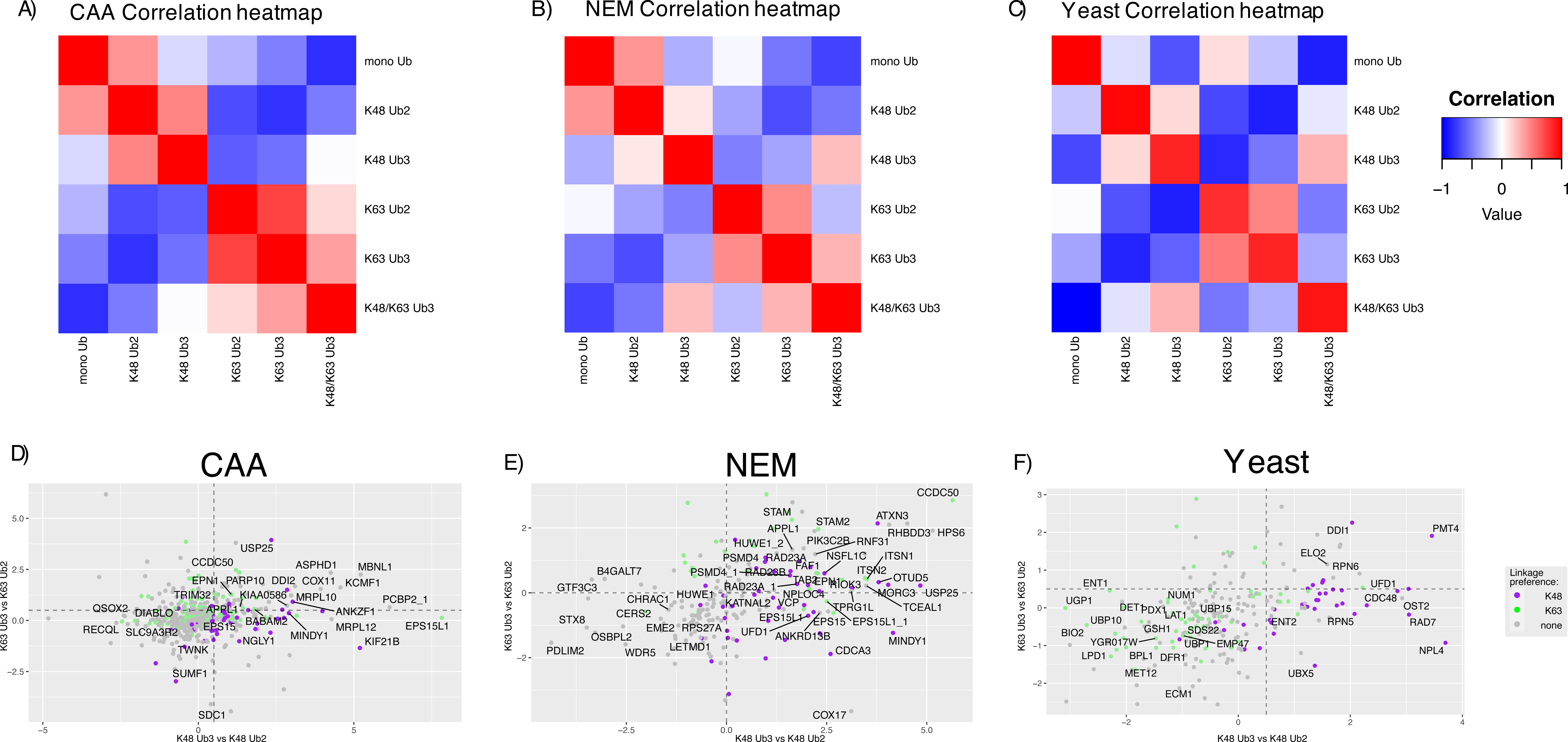
Ub interactome correlation and length-specific interactors. A, B and C) Correlation of Ub chain interactomes within Hela (A) CAA and (B) NEM, and yeast datasets. Spearman correlation calculated by comparing the moderated F value for each Ub pulldown of significant hits within each dataset. Significant hits identified by moderated F test of versus control filtered proteins. Adjusted P value cut off < 0.05. A and B) extended view of Figure 3 A and B. D, E and F) Scatterplots of Ub2 versus Ub3 comparison for both K48- and K63-linked Ub in D) CAA, E) NEM and F) yeast datasets (two-sample moderated T tests). Dot colours refer to linkage preference by two sample moderated T-test of K48-linked Ub3 versus K63-linked Ub3: purple is K48 significant, green is K63 significant and grey is not statistically significant in this comparison. Labelled proteins are significant in either Ub2 versus Ub3 comparison with D and F) Adj.P < 0.05 and E) AdjP < 0.005.

**Supplementary Figure 10:**
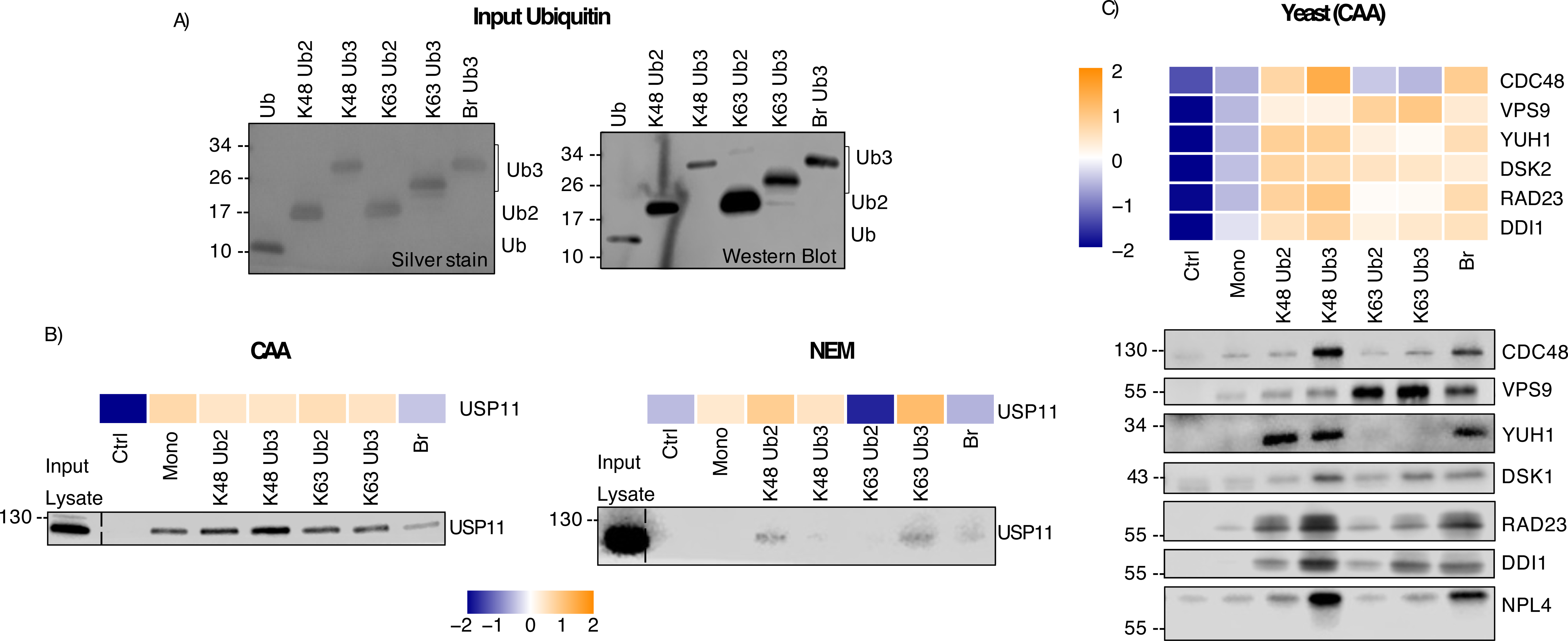
Western Blot validation of ubiquitin interactor enrichment. A) Input Ub chains by silver stain and Western blot with anti-Ub antibody. Experiment also shown in Figure 1B, 3E and F and 5A. B) Heatmap of MS-identified enrichment pattern combined with Western Blot of Ub enrichment of specific interactors from CAA- and NEM-treated lysate. Input and Ub pulldown seen in A) and Figure 1B. Blotted using anti-USP11 antibody. Heatmap generated using moderated F values for each bait type, before versus control filter. No P value cut off used. C) Heatmap of MS-identified enrichment pattern combined with Western blot of Ub enrichment of specific interactors from yeast lysate. Heatmap generated using moderated F values for each bait type, before versus control filtering. No P value cut off used. Blotted using anti-Cdc48, anti-Vps9, anti-YUH1, anti-Dsk2, anti-Rad23, anti-Ddi1 and anti-Npl4 antibodies.

**Supplementary Figure 11:**
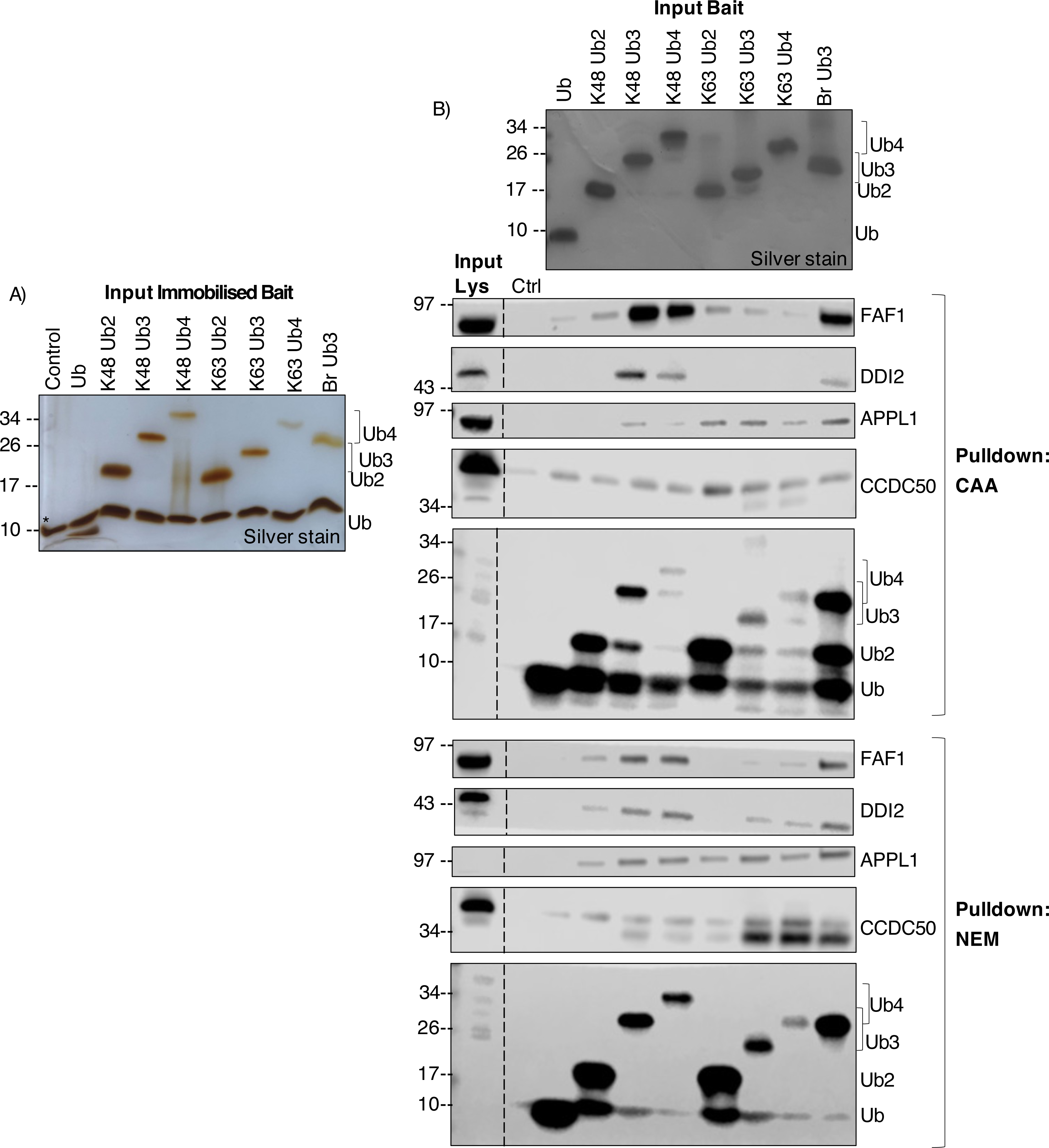
Western blot interactor enrichment including Ub4. A) Silver stain of streptavidin-immobilised Ub bait. * indicates streptavidin band from denaturation of streptavidin beads during sample prep for SDS-PAGE. B) Western Blot of Ub-interactor enrichment of MS-identified chain length-dependent interactors from CAA- and NEM-treated lysate. Blotted using anti-FAF1, anti-DDI2, anti-APPL1, anti-CCDC50 and anti-Ub antibodies. Blot from same experiment also shown in Figure 5B.

**Supplementary Figure 12:**
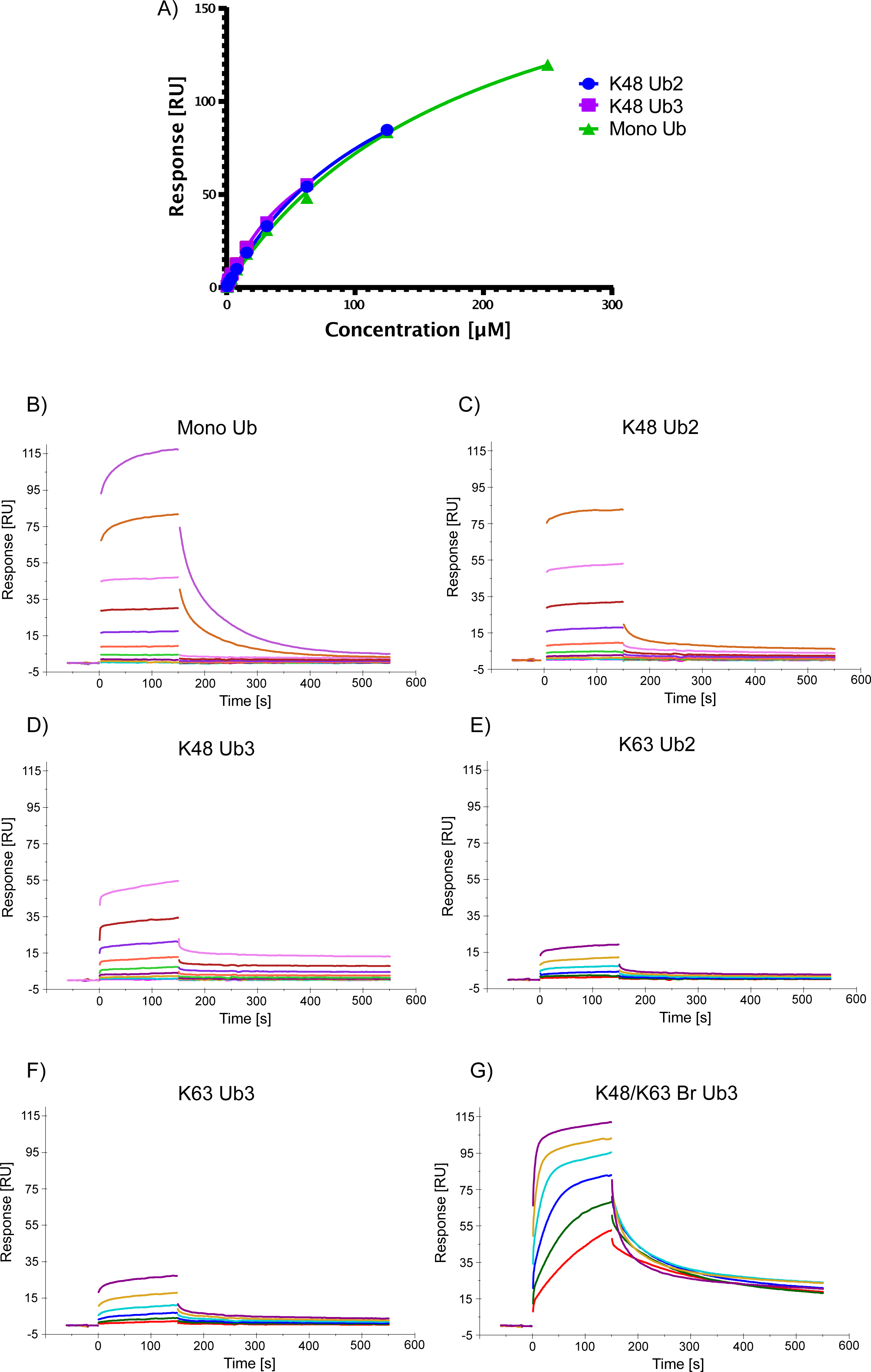
HIP1 Ub binding assay raw data. A) Overlayed affinity plots of triplicate measurements with determined K_D_ values and respective Standard Deviations (SD) of K48-linked Ub2, K48-linked Ub3 and mono Ub interacting with immobilized biotinylated HIP1 protein. Averaged response values [RU] at equilibrium were plotted against the injected concentration [µM] of respective analytes determined by SPR multi cycle format and fitted according to a steady state affinity model. B-F) Sensorgrams of (A) mono Ub, (B) K48 Ub2, (C) K48 Ub3, (D) K63 Ub2, (E) K63 Ub3 and (F) K48/K63 branched Ub3 binding to immobilized HIP1, by surface plasma resonance.

